# Quantitative proteomics identifies novel PIAS1 substrates involved in cell migration and motility

**DOI:** 10.1101/612622

**Authors:** Chongyang Li, Francis P. McManus, Cédric Plutoni, Cristina Mirela Pascariu, Trent Nelson, Lara Elis Alberici Delsin, Gregory Emery, Pierre Thibault

**Affiliations:** Institute for Research in Immunology and Cancer, Université de Montréal, Québec, Canada; Department of Molecular Biology, Université de Montréal, Québec, Canada; Department of Pathology and Cell Biology, Université de Montréal, Québec, Canada; Department of Chemistry, Université de Montréal, Québec, Canada; Department of Biochemistry, Université de Montréal, Québec, Canada

**Keywords:** SUMOylation, PIAS1, Vimentin, cell invasion, quantitative proteomics

## Abstract

The Protein Inhibitor of Activated STAT 1 (PIAS1) is an E3 SUMO ligase that plays important roles in various cellular pathways, including STAT signaling, p53 pathway, and the steroid hormone signaling pathway. PIAS1 can SUMOylate PML (at Lys-65 and Lys-160) and PML-RARα promoting their ubiquitin-mediated degradation. Increasing evidence shows that PIAS1 is overexpressed in various human malignancies, including prostate and lung cancers. To understand the mechanism of action of PIAS1, we developed a quantitative SUMO proteomic approach to identify potential substrates of PIAS1 in a system-wide manner. Our analyses enabled the profiling of 983 SUMO sites on 544 proteins, of which 204 SUMO sites on 123 proteins were identified as putative PIAS1 substrates. These substrates are involved in different cellular processes, such as transcriptional regulation, DNA binding and cytoskeleton dynamics. Further functional studies on Vimentin (VIM), a type III intermediate filament protein involved in cytoskeleton organization and cell motility, revealed that PIAS1 exerts its effects on cell migration and cell invasion through the SUMOylation of VIM at Lys-439 and Lys-445 residues. VIM SUMOylation was necessary for its dynamic disassembly, and cells expressing a non-SUMOylatable VIM mutant showed reduced levels of proliferation and migration. Our approach not only provides a novel strategy for the identification of E3 SUMO ligase substrates, but also yields valuable biological insights into the unsuspected role of PIAS1 and VIM SUMOylation on cell motility.

## INTRODUCTION

The small ubiquitin-like modifier (SUMO) protein is an ubiquitin-like (UBL) protein that is highly dynamic and can reversibly target lysine residues on a wide range of proteins involved in several essential cellular events, including protein translocation and degradation, mitotic chromosome segregation, DNA damage response, cell cycle progression, cell differentiation and apoptosis^1^. SUMO proteins are highly conserved through evolution, and the human genome encodes 4 SUMO genes, of which 3 genes (SUMO-1, SUMO-2 and SUMO-3) are ubiquitously expressed in all cells^1, 2^. Prior to conjugation, the immature SUMO proteins are C-terminally processed by sentrin-specific proteases (SENPs) ^3^. These proteases also cleave the isopeptide bond formed between the ε-amino group of the acceptor lysine residues and the C-terminus residue of the conjugated SUMO proteins. The conjugation of SUMO to target proteins requires an E1 activating enzyme (SAE1/2), an E2 conjugating enzyme (UBC9) and one of several E3 SUMO ligases^4^. Unlike ubiquitination, *in vitro* SUMOylation can occur without E3 SUMO ligases, although enhanced substrate specificity is conferred by E3 SUMO ligases ^5^. It is believed that SUMOylation events occurring without the aid of E3 SUMO ligases arise primarily on the consensus motif composed of ψKxE, where ψ represents a large hydrophobic residue and x, any amino acid^6^. To date, several structurally unrelated classes of proteins appear to act as E3 SUMO ligases in mammalian cells, such as the protein inhibitor of activated STAT (PIAS) family of proteins, Ran-binding protein 2 (RanBP2), the polycomb group protein (Pc2), and topoisomerase I- and p53-binding protein (TOPORS)^7, 8^.

PIAS orthologs can be found through eukaryote cells, and comprise four PIAS proteins (PIAS1, PIASx (PIAS2), PIAS3, and PIASy (PIAS4)) that share a high degree of sequence homology^9^. Overall, five different domains or motifs on PIAS family proteins recognize distinct sequences or conformations on target proteins, unique DNA structures, or specific “bridging” molecules to mediate their various functions^10^. An example of this is the PIAS Scaffold attachment factor (SAP) domain which has a strong affinity towards A–T rich DNA^11^ and binds to Matrix attachment regions DNA^12^, in addition to having an important role in substrate recognition^13^. The PINIT motif affects subcellular localization and contributes to substrate selectivity^14, 15^. The Siz/PIAS RING (SP-RING) domain interacts with UBC9 and facilitates the transfer of SUMO to the substrate^16^. The PIAS SIM (SUMO interaction motif) recognizes SUMO moieties of modified substrates and alters subnuclear targeting and/or assembly of transcription complex^16–18^. While several functions have been attributed to these domains, relatively little is known about the role of the poorly conserved C-terminus serine/threonine-rich region.

PIAS1 is one of the most well studied E3 SUMO ligases, and was initially reported as the inhibitor of signal transducers and activators of transcription 1 (STAT1)^19^. Previous studies indicated that PIAS1 interacts with activated STAT1 and suppresses its binding to DNA ^8^. PIAS1 overexpression was reported in several cancers, including prostate cancer, multiple myeloma, and B cell lymphomas^20–23^. PIAS1 can SUMOylate the Focal adhesion kinase (FAK) at Lys-152, a modification that dramatically increases its ability to autophosphorylate Thr-397, activate FAK, and promotes the recruitment of several enzymes including Src family kinases^24^. In yeast, Lys-164 SUMOylation on Proliferating Cell Nuclear Antigen (PCNA) is strictly dependent on the PIAS1 ortholog Siz1, and is recruited to the anti-recombinogenic helicase Srs2 during S-phase^25^. PIAS1 can also regulate oncogenic signaling through the SUMOylation of promyelocytic leukemia (PML) and its fusion product with the retinoic acid receptor alpha (PML-RARα) as observed in acute promyelocytic leukemia (APL)^26^. In addition to its regulatory role in PML/ PML-RARα oncogenic signaling, PIAS1 has been shown to be involved in the cancer therapeutic mechanism of arsenic trioxide (ATO). This is accomplished by ATO promoting the hyper-SUMOylation of PML-RARα in a PIAS1-dependent fashion, resulting in the ubiquitin-dependent proteasomal degradation of PML-RARα and APL remission^26^. In B cell lymphoma, PIAS1 has been reported as a mediator in lymphomagenesis through SUMOylation of MYC, a proto-oncogene transcription factor associated with several cancers. SUMOylation of MYC leads to a longer half-life and therefore an increase in oncogenic activity^23^. Altogether, these reports suggest that PIAS1 could promote cancer cell growth and progression by regulating the SUMOylation level on a pool of different substrates.

In this study, we first evaluate the effect of PIAS1 overexpression in HeLa cells. PIAS1 overexpression has a significant influence on cell proliferation, cell migration and cancer cell invasion. To identify putative PIAS1 substrates, we developed a system level approach based on quantitative SUMO proteomic analysis ^27^ to profile changes in protein SUMOylation in cells overexpressing this E3 SUMO ligase. Our findings revealed that 204 SUMO sites on 123 proteins were regulated by PIAS1. Bioinformatic analysis indicated that many PIAS1 substrates are involved in transcription regulation pathways and cytoskeleton organization. Interestingly, several PIAS1 substrates, including cytoskeletal proteins (Actin filaments, Intermediate filaments and Microtubules), were SUMOylated at lysine residues located in non-consensus motif. We confirmed the SUMOylation of several PIAS1 substrates using both *in vitro* and *in vivo* SUMOylation assays. Further functional studies revealed that PIAS1 mediated the SUMOylation of vimentin (VIM) at two conserved sites on its C-terminus that affect the dynamic disassembly of this intermediate filament protein.

## RESULTS

### PIAS1 Overexpression Promotes cell Proliferation and Motility

To investigate the physiological function of PIAS1 in HeLa cells, we overexpressed PIAS1 and evaluated its expression level by western blot. The abundance of PIAS1 in HeLa cells was increased by 2.5 fold at 48 h post-transfection (Fig. 1a). PIAS1 overexpression promotes HeLa cell proliferation by ∼50% (Fig. 1b). We further examined the phenotypic effects of PIAS1 overexpression on cell migration and cell invasion using wound-healing and Boyden chamber invasion assays. The migration and invasion ability of Hela cells were both increased after PIAS1 overexpression (Figs. 1c-f). Taken together, these results highlight the role that PIAS1 plays in regulating cell growth, cell cycle, cell migration and cell invasion of HeLa cells.

**Figure 1.**
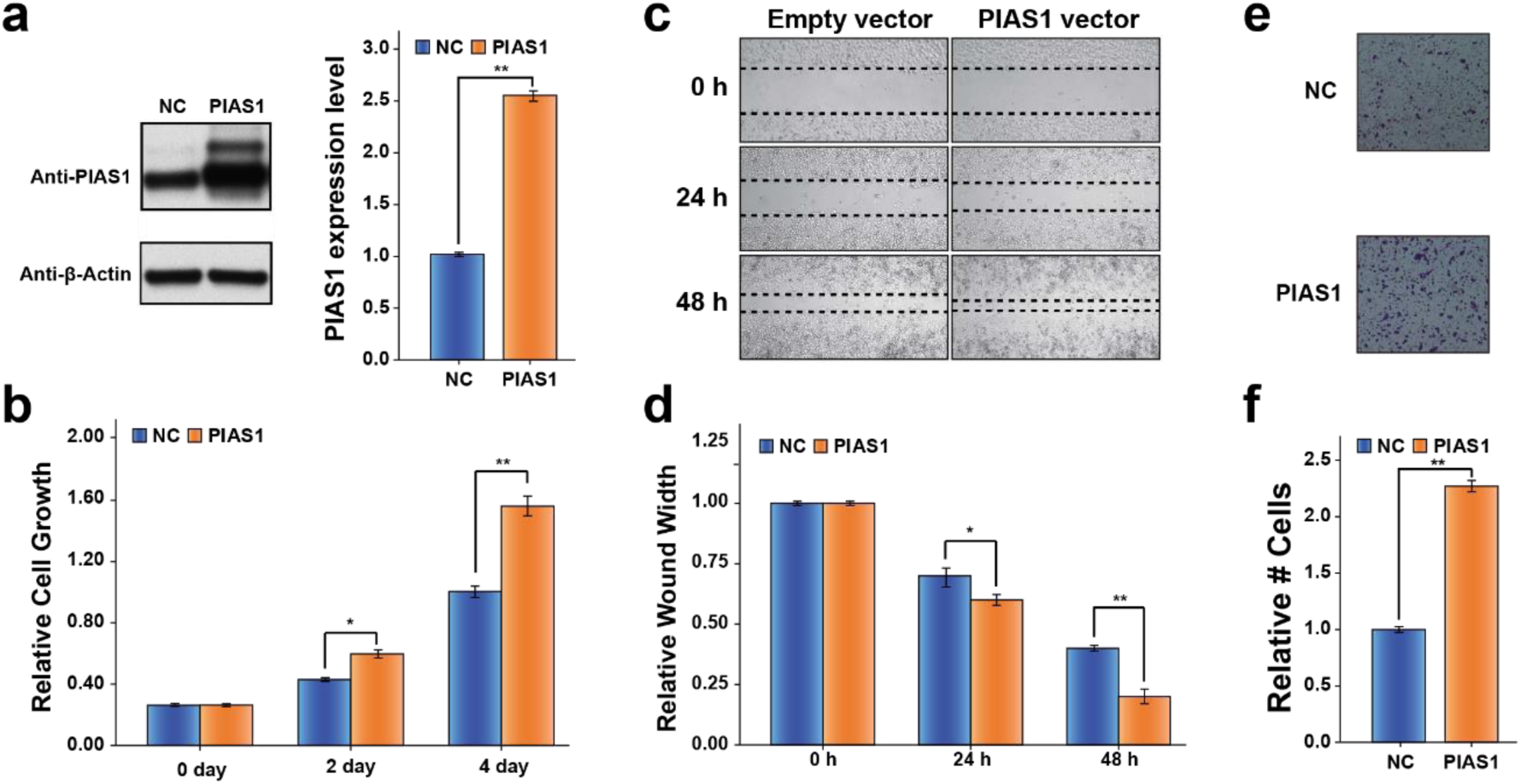
Functional effects of PIAS1 overexpression on HeLa cells. (a) HeLa cells were transfected with Myc-PIAS1 or Empty vector (NC) for 48 hr. PIAS1 overexpression efficiency was determined by western blot. Actin was used as a loading control. (b) PIAS1 overexpression in HeLa cells significantly promotes cell growth. (c) PIAS1 overexpression increases cell migration as determined by a wound-healing assay. (d) Quantification of the wound healing assay. (e) Representative results from a Boyden chamber assay showing improved invasion capabilities of PIAS1 overexpression in HeLa cells. (f) Quantification of the Boyden chamber assay. (*p<0.05, **p<0.01, Student’s t-test).

### Identification of PIAS1 Regulated SUMOylation Sites by Quantitative Proteomics

To gain a better understanding of the role that PIAS1 plays in cell cycle regulation, cell proliferation, invasion and motility, we devised a large-scale SUMO proteomic approach to identify PIAS1 substrates in a site-specific manner (Fig. 2). We combined a SUMO remnant immunoaffinity strategy ^27^ with metabolic labeling (stable isotope labeling of amino acid in cell culture, SILAC) to study the global changes in protein SUMOylation upon PIAS1 overexpression. HEK293 cells stably expressing SUMO3m (Supplementary Fig. 1) were grown at 37 °C in media containing light (^0^Lys, ^0^Arg), medium (^4^Lys, ^6^Arg), or heavy (^8^Lys, ^10^Arg) isotopic forms of lysine and arginine. Three biological replicates were performed, and for each replicate, one SILAC channel was transfected with an empty vector while the other two were transfected with Myc-PIAS1 vectors (Fig. 2a). At 48 h post-transfection, an equal amount of cells from each SILAC channel were harvested and combined before lysis in a highly denaturant buffer. PIAS1 overexpression efficiency in HEK293 SUMO3m cells was evaluated by western blot (Fig. 2b). Protein extracts were first purified by Ni-NTA beads to enrich SUMO-modified proteins and digested on beads with trypsin (Fig. 2c). Following tryptic digestion, SUMO-modified peptides were immunopurified using an antibody directed against the NQTGG remnant that is revealed on the SUMOylated lysine residue. Next, peptides were fractionated by offline strong cation exchange (SCX) STAGE tips and analyzed by LC-MS/MS on a Tribrid Fusion instrument. To determine that abundance changes were attributed to SUMOylation and not to change in protein expression, we also performed quantitative proteomic analyses on the total cell extracts from PIAS1 overexpression (Fig. 3a and Supplementary Fig. 2). PIAS1 overexpression caused a global increase in protein SUMOylation with negligible changes on protein abundance (Fig. 3b). In total, 12080 peptides on 1756 proteins (Fig. 3a, Supplementary Table 1) and 983 SUMO peptides on 544 SUMO proteins (Fig. 3b, Supplementary Table 2) were quantified for the proteome and SUMO proteome analyses, respectively. A total of 204 SUMOylation sites on 123 proteins were found to be upregulated by PIAS1 overexpression including known substrates such as PCNA and PML. A summary of these analyses is shown in Fig. 3 c.

**Figure 2.**
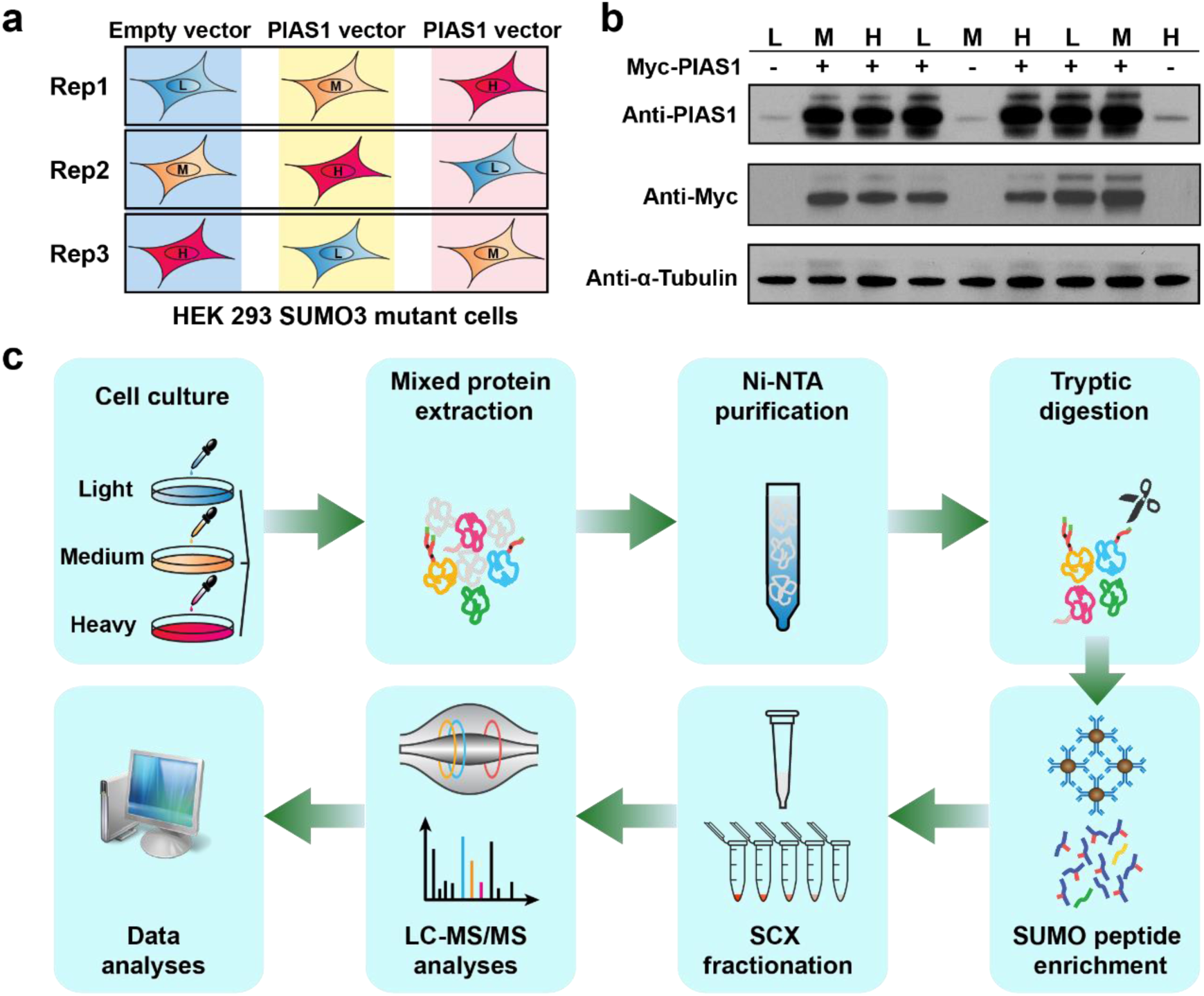
Workflow for the identification of PIAS1 substrates. (a) HEK293 SUMO3m cells were cultured in SILAC medium with reverse labeling in biological triplicates. For replicate 1, PIAS1 overexpression was performed in medium and heavy channels with Myc-PIAS1 vector, while the cells cultured in light media were transfected with the pcDNA3.0 (Empty vector). For replicates 2 and 3 the empty vector was transduced in the medium and heavy labelled cells, respectively. (b) Western blot showing the level of overexpression of PIAS1 in the transfected cells. Detection of PIAS1 overexpression by both Anti-PIAS1 antibody and Anti-Myc antibody. β-Tubulin is used as loading control. (c) SILAC labelled cells were lysed and combined in a 1:1:1 ratio based on protein content. SUMOylated proteins were enriched from the cell extract on an IMAC column prior to their tryptic digestion. After desalting and drying, peptides containing the SUMO3 remnant were enriched using a custom anti-K-ε-NQTGG antibody that was crosslinked on magnetic beads [4]. Enriched peptides were further fractionated on SCX columns and injected on a Tribrid Fusion mass spectrometer. Peptide identification and quantification were performed using MaxQuant.

**Figure 3.**
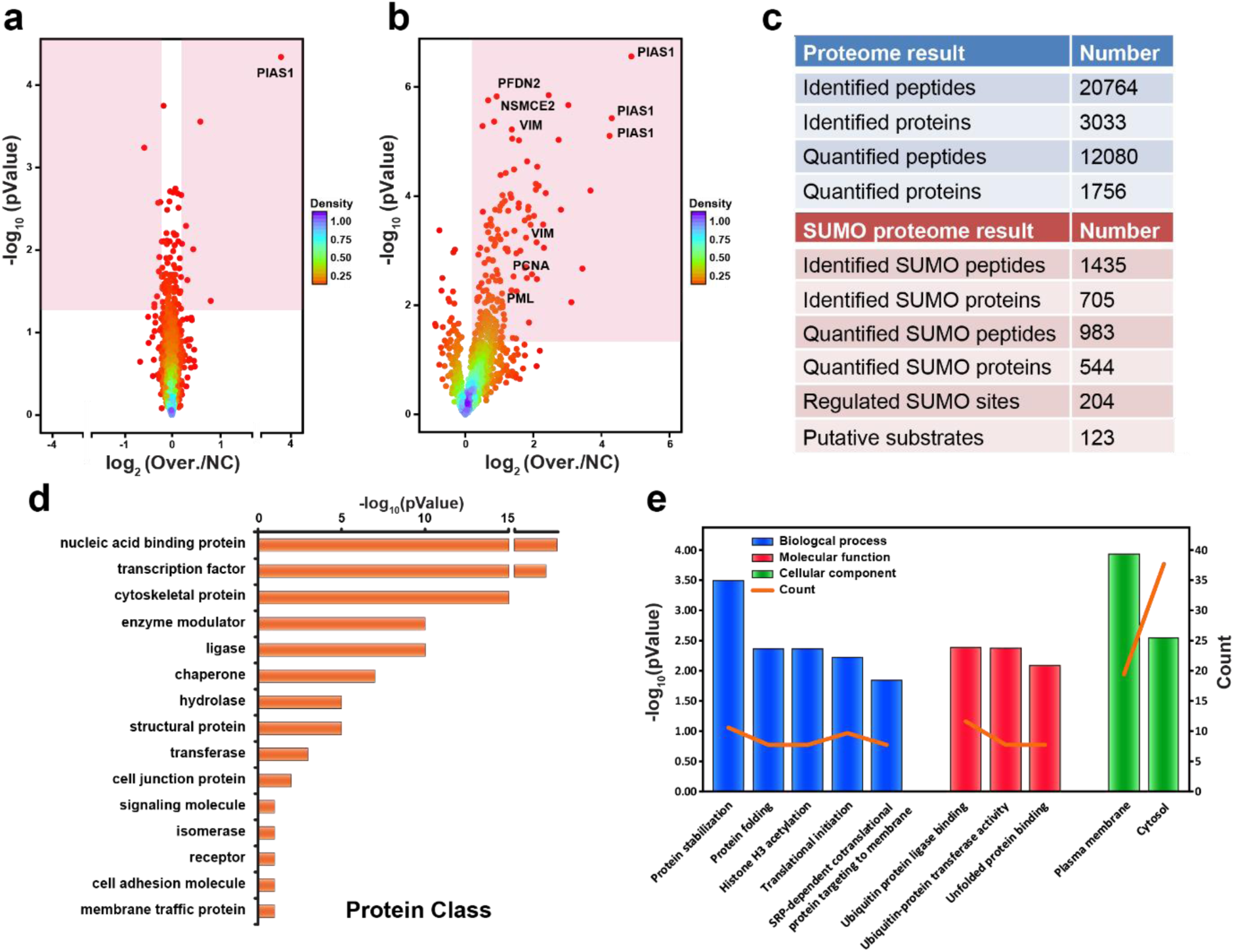
Statistical analyses of mass spectrometry results and bioinformatics analyses of identified PIAS1 substrates. (a) Volcano plots showing the global proteome changes in cells overexpressing PIAS1 (Over.) vs. control cells (NC). Individual proteins are represented by points. The area of the volcano plot where protein abundance changes are significantly regulated (p-value of <0.05) are shaded in pink. (b) Volcano plots showing the global SUMOylation changes in cells overexpressing PIAS1 (Over.) vs. control cells (NC). Individual SUMOylation sites are represented by points. The area of the volcano plot where SUMO sites are significantly up-regulated (p-value of <0.05) is shaded in pink. (c) Summary of identified and quantified peptides and proteins in both the proteome and SUMOylome experiments. (d) Functional classification of PIAS1 substrates using PANTHER (Protein Analysis Through Evolutionary Relationships) (http://www.pantherdb.org). (e) GO term enrichment distribution of the identified PIAS1 substrates using DAVID 6.8 (https://david.ncifcrf.gov/).

Protein classification ontology analysis of the PIAS1 substrates using PANTHER clustered the targets into 15 groups (Fig. 3d). PIAS1 mediated SUMOylation predominantly occurred on nucleic acid binding proteins, transcription factors, cytoskeletal proteins, enzyme modulators, ligases and chaperone proteins. We next classified putative PIAS1 substrates by their gene ontology (GO) molecular function, biological process and cellular components (Fig. 3e) using the whole identified SUMOylome as background. GO cellular component classification revealed that PIAS1 substrates were enriched in the plasma membrane and cytosol compared to the global SUMOylome (Fig. 3e). This suggests that PIAS1 substrates may undergo nucleocytosolic shuttling upon SUMOylation. GO biological process analysis revealed that identified PIAS1 substrates are involved in a variety of biological processes, such as protein stabilization, protein folding, histone H3 acetylation, translational initiation, and signal recognition particle (SRP)-dependent co-translational protein targeting to membrane (Fig. 3e). GO molecular function analysis indicated that PIAS1 substrates are associated with ubiquitin protein ligase binding, ubiquitin-protein transferase activity and unfolded protein binding (Fig. 3e). Indeed, PIAS1 regulates the SUMOylation of several proteins whose roles in the cell are diverse. Much like global SUMOylation, PIAS1 mediated SUMOylation may play a role in several biological processes that are independent from each other.

Previous SUMO proteome analyses indicated that under unstressed conditions, approximately half of acceptor lysine residues are found in the SUMO consensus and reverse consensus motifs^28^. Since it is believed that SUMOylation that occurs on the consensus motif does not absolutely require an E3 ligase, we surmised that PIAS1 mediated SUMOylation may occur at non-consensus motifs. We therefore compared the amino acid residues surrounding the SUMOylation sites that are regulated by PIAS1 to those of the whole SUMO proteome (Supplementary Fig. 3a). As anticipated, the sequences surrounding the PIAS1 mediated SUMOylation sites are depleted in glutamic acid at position +2 and depleted of large hydrophobic amino acids at position −1, consistent with the reduction of the consensus sequence. Indeed, E3 SUMO ligases appear to aid in the SUMOylation of lysine residues that reside in non-canonical regions. Furthermore, we investigated the local secondary structures and solvent accessibility of PIAS1 substrates surrounding SUMO sites using NetSurfP-1.1 software (Supplementary Fig. 3b). We observed that one-third of PIAS1 regulated SUMOylation sites are located within α-helix and approximately one-tenth within β-strand. In contrast, the majority of SUMOylated lysine residues in the SUMO proteome are localized in coil regions. Taken together, these results support the notion that PIAS1 mediated SUMOylation preferentially occurs on structured regions of the protein, which may help substrate recognition by PIAS1. Additionally, we noted that PIAS1 mediated SUMOylation occurred primarily on solvent exposed lysine residues, which was also the case for the global SUMOylome. These results suggest that PIAS1 may not impart conformational changes to its substrate upon binding since it does not promote SUMOylation on lysine residues that would otherwise be buried within the core of the substrate. Overall, PIAS1 promotes the ability of UBC9 to SUMOylate lysine residues that are present in non-consensus sequences located on ordered structures of the proteins.

To better understand the cellular processes regulated by PIAS1, a STRING analysis was performed to analyze the interaction network of putative PIAS1 substrates. This network highlights the presence of highly connected interactors from promyelocytic leukemia protein (PML) nuclear body, transcriptional factors, cytoskeletal proteins and RNA binding proteins (Fig. 4). PIAS1 was previously shown to colocalize to PML nuclear body and to regulate oncogenic signaling through SUMOylation of PML and its gene translocation product PML-RARα associated with acute promyelocytic leukemia (APL) ^26^.

**Figure 4.**
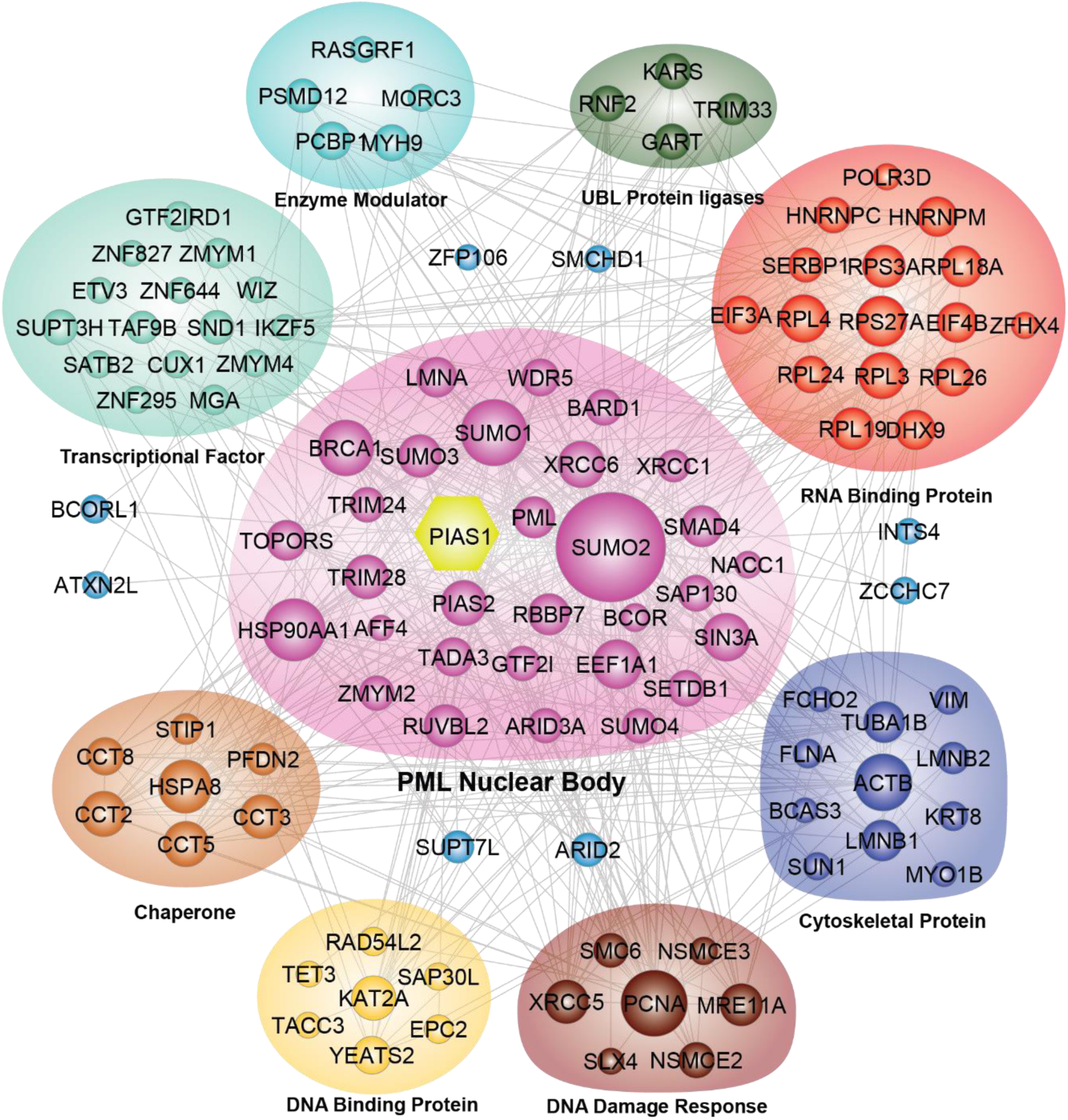
Protein-Protein Interaction Network of PIAS1 substrates. STRING network of PIAS1 substrates and their interacting partners. Proteins are grouped according to their GO terms.

Interestingly, we also found that several putative PIAS1 substrates were associated with cytoskeletal organization including β-actin (ACTB), α-tubulin (TUBA1B), vimentin (VIM) and Keratin 8 (KRT8), in addition to several other intermediate filament proteins. Of note, Keratin 8 was reported to be SUMOylated *in vitro* and *in vivo*, a modification that changes its dynamic assembly and is increased in cells and tissues during apoptosis, oxidative stress, and phosphatase inhibition^29^. The actin filaments, intermediate filaments, and microtubules that form the cytoskeleton of eukaryotic cells are responsible for cell motility and division. They also help establish cell polarity, which is required for cellular homeostasis and survival^30^. Moreover, one SUMO site on β-actin (ACTB) and five SUMO sites on α-tubulin (TUBA1B) were found at non-consensus motif regions, indicating an important role for PIAS1 in substrate protein dynamics during SUMOylation. Additionally, we evaluated the degree of evolutionary conservation of these modified lysine residues. Surprisingly, all SUMOylated lysine residues analyzed are highly conserved across different species (Supplementary Figs. 4 and 5). In our data, we found intermediate filaments (IFs) to be major targets of PIAS1 among cytoskeletal proteins. Our results highlight that PIAS1 mediates the SUMOylation of the type II IF keratin 8 protein at Lys-421 and the type III IF VIM protein at Lys-439 and Lys-445, both on their respective tail domains. Moreover, PIAS1 promotes the SUMOylation of several type V IF proteins (e.g. Prelamin A/C, Lamins B1 and B2) on their Rod domains (Supplementary Fig. 6).

**Figure 5.**
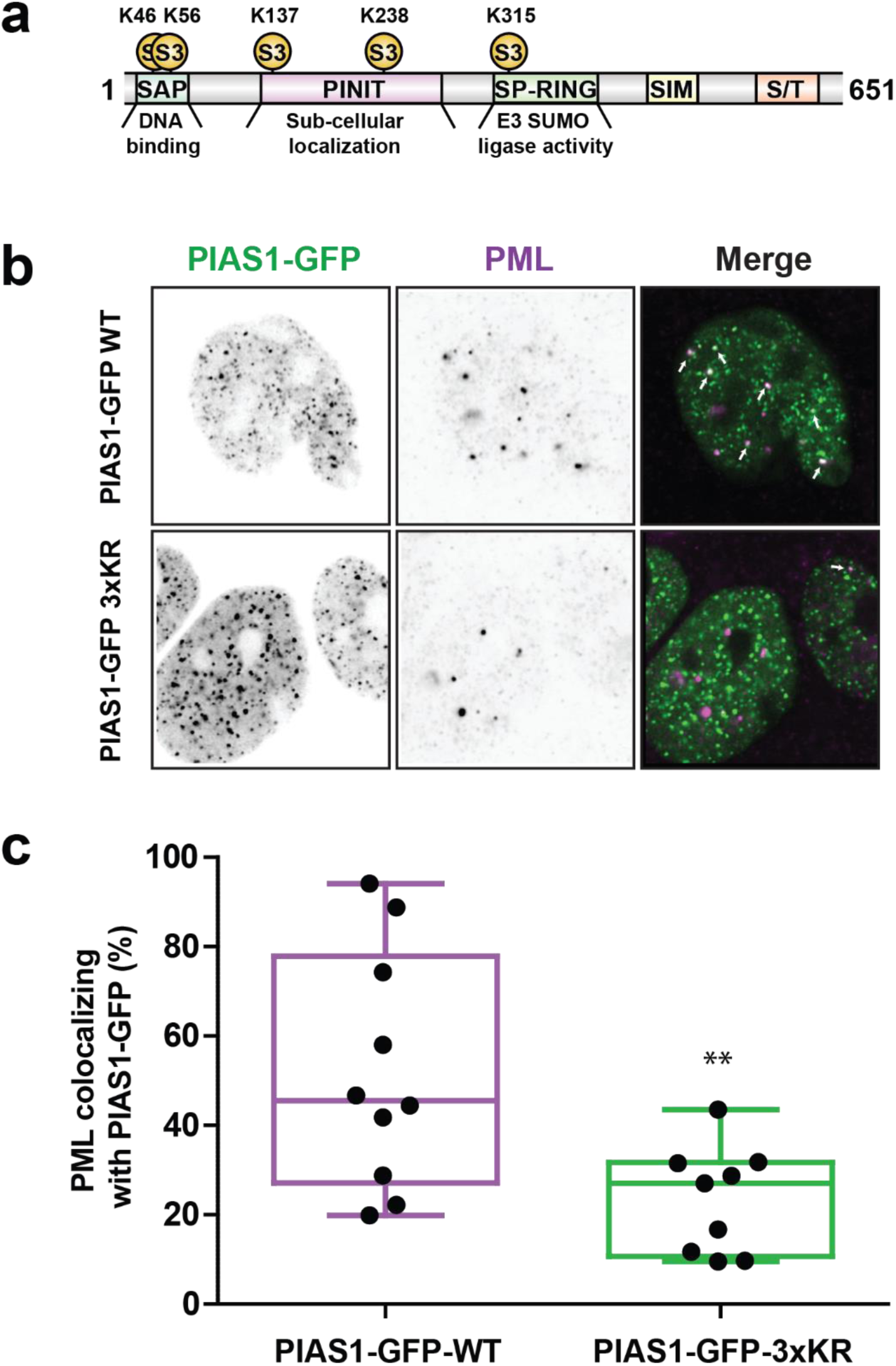
SUMOylation of PIAS1 promotes its PML localization. (a) Distribution of SUMO sites identified on PIAS1. Three SUMO sites (K137, K237 and K315) were identified in the dataset. K137 and K238 are located in PINIT domain while K315 is located in SP-RING domain. (b) HEK293 SUMOm cells were co-transfected with PIAS1-GFP-WT or PIAS1-GFP-3xKR and immunofluorescence was performed with anti-PML (scale bar, 10 μm). (c) Box plot graph showing the ratio of PIAS1-PML co-localization relative to the total cell signal. (**p<0.01, Student’s t-test, n=10 cells/condition)

### Validation of identified PIAS1 substrates by in vitro and in vivo SUMOylation assays

Next, we selected E3 SUMO-protein ligase NSE2 (NSMCE2), prefoldin subunit 2 (PFDN2) and VIM, which were identified in SUMO proteomic experiments as putative PIAS1 substrates for further validation. We performed both *in vitro* and *in vivo* SUMOylation assays to confirm that these sites were regulated by PIAS1. For the *in vitro* SUMO assay, we incubated individual SUMO substrates with SUMO-activating E1 enzyme (SAE1/SAE2), UBC9, SUMO-3 with or without PIAS1 in the presence of ATP. We also used PCNA, a known PIAS1 substrate, as a positive control. After 4h incubation at 37 °C, the western blots of each substrate showed either single or multiple bands of higher molecular weight confirming the SUMOylated products. Separate LC-MS/MS experiments performed on the tryptic digests of the *in vitro* reactions confirmed the SUMOylation of NSMCE2 at residues Lys-90, Lys-107, and Lys-125, and PFDN2 at residues Lys-94, Lys-111, Lys-132, and Lys-136. While UBC9 alone can SUMOylate these substrates, we noted an increasing abundance of SUMOylated proteins when PIAS1 was present, confirming that the E3 SUMO ligase enhanced the efficiency of the conjugation reaction (Figure S7a). Interestingly, several SUMOylation sites that were regulated by PIAS1 on both NSMCE2 and PFDN2 were not located within SUMO consensus motifs, further supporting the motif analysis of the large-scale proteomic data (Supplementary Fig. 3a).

Furthermore, we examined whether PIAS1 contributes to substrate SUMOylation *in vivo*. HEK293-SUMO3 cells were co-transfected with Flag-NSMCE2, PFDN2 or VIM and Myc-PIAS1. Co-transfected cells were subjected to immunoprecipitation with anti-Flag agarose gel, followed by western blot with an anti-His antibody. The SUMOylation of substrates was minimally detected when only transfecting Flag-substrates. In contrast, overexpression of PIAS1 under the same experimental conditions led to a marked increase in the SUMOylation of these substrates (Supplementary Fig. 7b). These results further confirm our quantitative SUMO proteomics data and validate the proteins NSMCE2, PFDN2 and VIM as *bona fide* PIAS1 substrates.

### PIAS1 SUMOylation Promotes its Localization to PML-NBs

Interestingly, our large-scale SUMO proteomic analysis identified five SUMOylation sites on PIAS1 (Fig. 5a). Two of these sites (Lys-46, Lys-56) are located in the SAP domain, and may regulate the interaction of PIAS1 with DNA ^15^. We also identified two SUMOylated residues (Lys-137 and Lys-238) located within the PINIT domain of PIAS1, potentially affecting its subcellular localization^14^. The last SUMOylated site (Lys-315) of PIAS1 and is located next to an SP-RING domain, which may alter the ligation activity of PIAS1 ^31^. Of note, PIAS1 contains a SUMO interaction motif (SIM), and previous reports indicated that this ligase can localize to PML nuclear bodies in a SIM-dependent manner with SUMOylated PML^32^. As PML also contains a SIM motif, we were interested in three out of the five SUMO sites on PIAS1: Lys-137, Lys-238 and Lys-315. Since PIAS1 is SUMOylated at several sites and PML contains a SIM, we surmised that reciprocal interactions could be mediated through SUMO-SIM binding. Accordingly, we constructed a PIAS1-GFP vector and used site-directed mutagenesis to specifically mutate the PIAS1 lysine residues that are SUMO modified and are located within regions of PIAS1 that could interact with PML. As the SAP domain of PIAS1 is exclusively reserved for DNA binding, we excluded the SUMO-modified lysine residues in this domain when creating the mutant construct as it may affect its localization in a PML independent fashion. We therefore created the variant constructs by mutating the codons for Lys-K137, Lys-238 and Lys-315 to arginine codons. Several mutant genes were created, including single mutants of each site, double mutant combinations, and the triple mutant (PIAS1-GFP 3xKR). These mutant vectors were transfected into HEK293 SUMO3m cells and used to study the effects of SUMOylation at the various lysine residues on the PIAS1-PML colocalization. As evidenced by the immunofluorescence studies, approximately 45% of WT PIAS1-GFP colocalized with PML (Fig. 5b). No significant changes in the colocalization of PIAS1 and PML were observed when experiments were repeated using either single or double PIAS1-GFP mutants (data not shown). However, we noted a 50% reduction in PIAS1-GFP–PML colocalization when all three sites were mutated, suggesting a possible functional redundancy among these sites (Figs. 5b and 5c). The fact that the co-localization of PML and PIAS1-GFP was not totally abrogated for the triple mutant might be explained by residual interactions between SUMOylated PML and the SIM of PIAS1-GFP^32^. Taken together, these experiments confirmed that colocalization of PIAS1 at PML nuclear bodies is partly mediated by the SUMOylation of PIAS1 at Lys-137, Lys-238 and Lys-315 residues.

### VIM SUMOylation Promotes Cell Proliferation and Motility and Increases VIF Solubility

VIM is predominantly found in various mesenchymal origins and epithelial cell lines^33–35^. Increasing evidence shows that VIM plays key roles in cell proliferation^36^, migration^37^ and contractility^38^. Our data shows that two SUMO sites on VIM are regulated by PIAS1, both of which are located on the tail domain and are highly conserved across different species (Fig. 6a). To further investigate the function of PIAS1 mediated SUMOylation of VIM, we expressed a Flag-tagged VIM K439/445R double mutant (VIM^mt^) that is refractory to SUMOylation in HeLa cells, and compared the functional effects to cell expressing the wild-type Flag-tagged VIM (VIM^wt^). We transfected Flag-VIM^wt^ and Flag-VIM^mt^ into HeLa cells and used the empty Flag vector as a negative control. At 48 h post-transfection, the cells were harvested, lysed in 8 M urea and protein pellets were separated on SDS-PAGE. The ensuing western blot results show that protein abundance between VIM^wt^ and VIM^mt^ are similar; yet, the SUMO level on VIM^mt^ is undetectable (Fig. 6b). Growth assays demonstrate that VIM^wt^ expression has no effect on HeLa cell proliferation while VIM^mt^ expression results in a significant inhibition of cell growth (Fig. 6c). Moreover, we examined the effect of VIM^wt^ and VIM^mt^ expression on cell migration and invasion using wound-healing and Boyden chamber invasion assays. VIM^wt^ significantly promotes cell migration and invasion, which is in line with the results obtained in HepG2 cells^39^. However, VIM^mt^ conferred no effect on cell migration or invasion (Figs. 6d-g). These results indicate that SUMOylation of VIM plays a role in cell growth, migration and invasion, presumably by regulating VIM IF function and/or formation.

**Figure 6.**
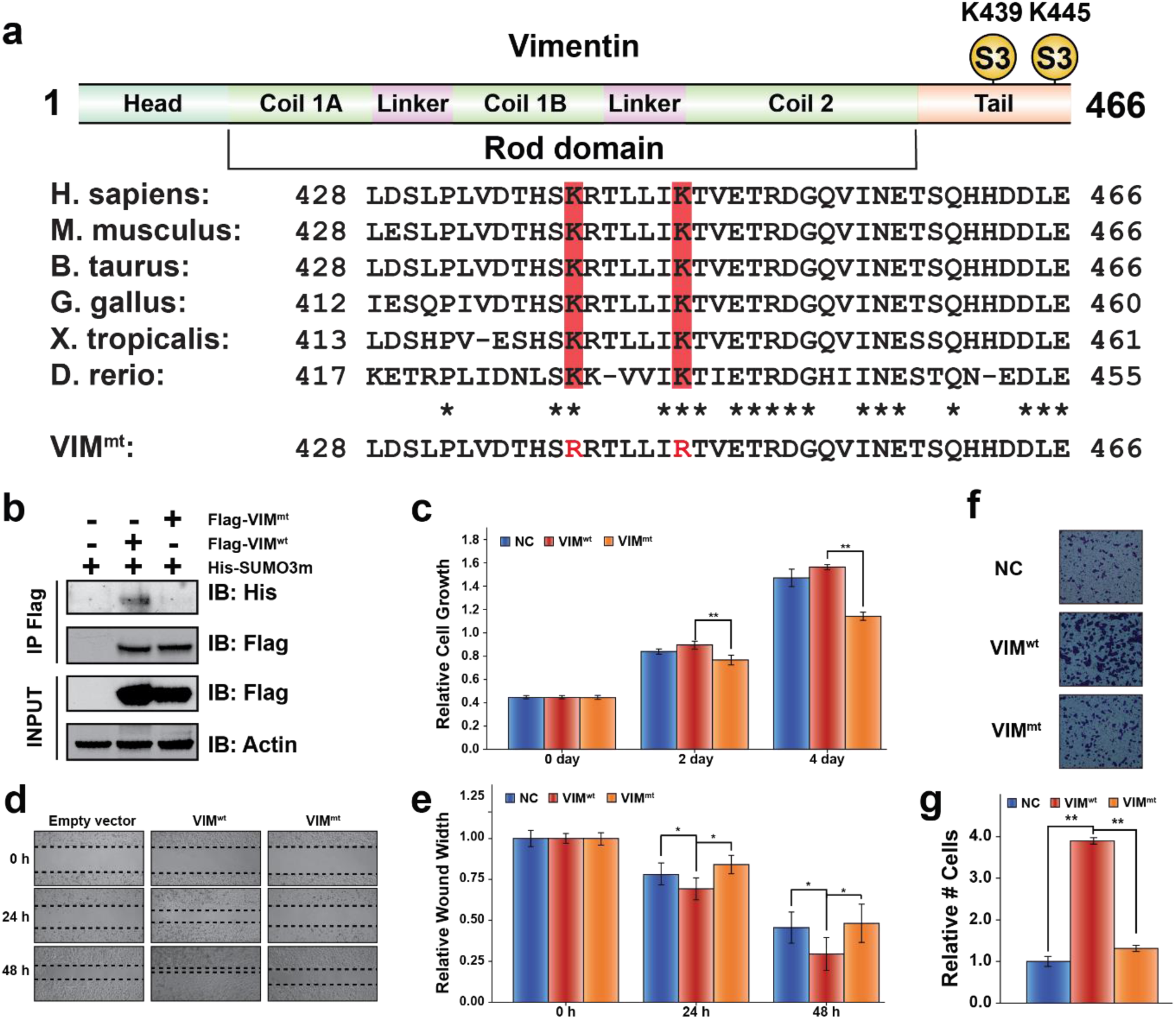
SUMOylation of Vimentin (VIM) is required for proper cell growth, migration and invasion in HeLa cells. (a) VIM sequence indicating SUMOylation sites at Lys 439 and Lys 445 localized in the tail domain, and highly conserved across six different species. (b) Western blot analysis of HeLa cells transfected with an empty vector as a negative control, Wild-type vimentin (Flag-VIM^wt^), and Vimentin K439, 445R, double mutant (Flag-VIM^mt^), showing the absence of SUMOylation on the Flag-VIM^mt^ protein. (c) VIM^mt^ expression inhibits proper cell growth. (d) VIM^mt^ expression inhibits proper cell migration as determined by a wound-healing assay. (e) Quantification of the wound healing assay. (f) Effect of VIM^wt^ and VIM^mt^ overexpression on cell invasion, as determined in a Boyden chamber assay. (g) VIM^mt^ expression significantly inhibits HeLa cell invasion. (*p<0.05, **p<0.01, Student’s t-test).

Next, we investigated the function of VIM SUMOylation on the dynamics assembly of IFs. Both Keratin and lamin A IFs formation and solubility have been reported to be regulated by SUMOylation, while such properties have yet to be uncovered for VIM IFs^40^. For example, the SUMOylation of lamin A at Lys-201, which is found in the highly conserved rod domain of the protein, results in its proper nuclear localization^41^. Unlike lamin A SUMOylation, keratin SUMOylation is not detected under basal conditions. However, stress-induced keratin SUMOylation has been observed in mouse and human in chronic liver injuries. Additionally, keratin monoSUMOylation is believed to increase its solubility, while hyperSUMOylation promotes its precipitation^42^.

We surmised that SUMOylation on VIM would alter its solubility, akin to the properties observed for keratin. Accordingly, we transfected an empty vector as a negative control, VIM^wt^ and VIM^mt^ into HeLa cells and lysed the cells with a RIPA buffer. Samples were fractionated into RIPA soluble and insoluble fractions. Western blot analysis of these samples shows that VIM^wt^ is preferentially located in the RIPA soluble fraction, while VIM^mt^ resides more in the insoluble fraction (Fig. 7a). Clearly, SUMOylation of VIM drastically increases its solubility.

**Figure 7.**
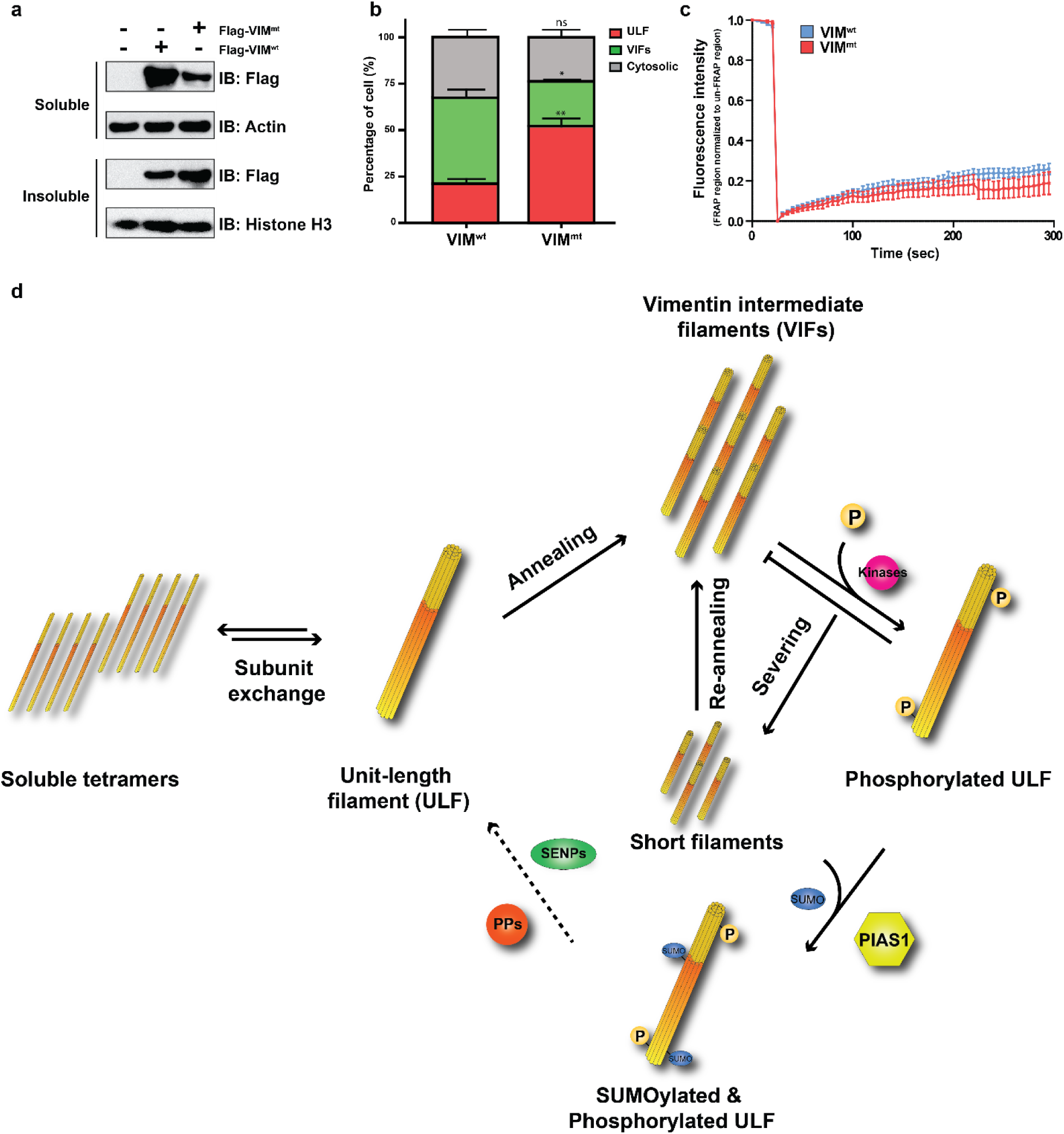
SUMOylation of Vimentin regulates its dynamic assembly. (a) HeLa cells were transfected with an empty vector as a negative control, Flag-VIM^wt^ and Flag-VIM^mt^ and separated into RIPA soluble and insoluble fractions. VIM protein levels were examined by western blot. (b) MCF-7 cells were transfected with Emerald-wild-type vimentin (VIM^wt^), and Emerald-vimentin K439, 445R, double mutant (VIM^mt^), and the proportion of unit-length filament (ULF), vimentin intermediate filament (VIF) and cytosolic vimentin between VIM^wt^ and VIM^mt^ was calculated under microscope (scale bar, 10 μm).(c) Fluorescence Recovery After Photobleaching (FRAP) assay of Emerald-wild-type vimentin (VIM^wt^), and Emerald-vimentin K439, 445R, double mutant (VIM^mt^) in MCF-7 cells. Line plot shows average fluorescence at each time point ± s.e.m. Differences between values for VIM^wt^ and VIM^mt^ were not statistically significant at all time points by Student’s t-test. (d) Model of the VIM dynamic assembly and disassembly. Vimentin is maintained in equilibrium between Unit-length Filament (ULF) and soluble tetramers. (Subunit exchange step). Vimentin filaments elongate by end-to-end annealing of ULF to form mature vimentin intermediate filament (VIF; annealing step). The phosphorylation-dependent shortening of VIF (severing step) involves phosphorylation on Ser 39 and Ser 56 of vimentin by several kinases, including Akt1. The short filaments can reanneal with another ULF to form new VIF (re-annealing step). However, the phosphorylated ULF are not amenable to the re-annealing process. These phosphorylated ULF products are subject to PIAS1-mediated SUMOylation, stimulating the dephosphorylation of the phosphorylated ULF, and subsequently reenter to either subunit exchange process or VIF maturation process.

IF networks are subject to regulation by a number of phosphatases and kinases^43^. Stable VIM filaments undergo extensive reorganization upon phosphorylation, a posttranslational modification that controls VIM assembly and disassembly^44–46^. Multiple phosphorylation sites have now been identified on VIM^47^. Those that induce disassembly are located in the N-terminal head domain and are phosphorylated by multiple protein kinases. Phosphorylation of the head region increases the distance between the two head groups of the VIM dimer, thus rendering the VIM tetramer incapable of assembling into a filament. Activation of Akt in soft-tissue sarcoma (STS) cells promotes the interaction of the head region of VIM and the tail region of Akt, resulting in the phosphorylation of VIM at Ser-39, further enhancing cell motility and invasion^41^.

Next, we sought to examine if VIM SUMOylation alters its phosphorylation status, which could lead to changes in VIM IF dynamic assembly/disassembly. Accordingly, we separated protein extracts from VIM^wt^ and VIM^mt^ by SDS-PAGE, excised bands that corresponded to the soluble and insoluble VIM, and performed in-gel trypsin digestion followed by LC-MS/MS (Supplementary Fig. 8). We identified several phosphorylated serine residues and one phosphorylated threonine residue on VIM (Supplementary Table 3), which have also been reported in the literature (S5, S7, T20, S22, S26, S29, S39, S42, S51, S56, S66, S72, S73, S83, S226, T258 and S459). All sites except those located on the C-terminal were found to be hyperphosphorylated in the insoluble VIM^mt^ compared to the wild-type counterpart. Interestingly, we observed an increase phosphorylation of S39 in the insoluble pellet of VIM^mt^, a site known to be phosphorylated by Akt ^41^. The observation that VIM is hyperphosphorylated at its N-terminus in VIM^mt^ suggests a possible cross-talk between SUMOylation and phosphorylation.

To further understand how SUMOylation affects VIM dynamics *in vivo*, we transfected Emerald-VIM^wt^ or -VIM^mt^ vectors in VIM null MCF-7 cells. VIM null cells were employed to eliminate the contribution of endogenous VIM on the VIM dynamics, which could mask the phenotypic effects of our mutant. Using fluorescence microscopy we quantified the proportion of the various VIM structures for the two different Emerald tagged constructs. We found three major forms of VIM in the cells, which in accordance with the literature, were categorized as cytosolic, unit-length filament (ULF) and VIM intermediate filaments (VIFs) (Supplementary Fig. 9). Statistical analysis of the proportion of the VIM structures revealed that the SUMO conjugation deficient VIM (VIM^mt^) promoted the formation of ULF with a concomitant reduction in VIF formation compared to its wild-type counterpart (Fig. 7b). Taken together these results indicate that SUMOylation of VIM promotes the formation of VIFs from the ULF building blocks.

Although VIM filament growth primarily relies on elongation by the longitudinal annealing of ULFs via end-to-end fusion, recent studies suggest that the subunit exchange of tetramers within these filaments does occur^48^. To determine if the SUMOylation of VIM affects the subunit exchange rate of these filaments we performed fluorescence recovery after photobleaching (FRAP) assays (Supplementary Fig. 10). We monitored the recovery time of filament fluorescence up to 300 s after bleaching for both Emerald-VIM^wt^ and Emerald-VIM^mt^, and noted no statistical difference in recovery between constructs (Fig. 7c). This observation suggests that the SUMOylation of VIM is not involved in the subunit exchange of tetramers within filaments.

The model described in Fig. 7d combines the results from the proteomic, immunofluorescence and FRAP assays and describes the molecular mechanism of PIAS1 mediated control of VIM dynamics. Under physiological conditions, vimentin is maintained in equilibrium between ULF and soluble tetramers. The formation of VIM ULF has been shown to occur spontaneously on the order of seconds *in vitro* showing that this arrangement is thermodynamically favorable and proceeds rapidly without the need for protein modifications^49^. Vimentin filaments elongate by end-to-end annealing of ULFs and eventually form mature vimentin intermediate filaments (VIFs). This is followed by the breakdown of VIFs by severing, which involves phosphorylation on several residues found on the N-term of VIM^47, 50^. We show in this work that this hyper-phosphorylation occurs exclusively on the N-terminal of VIM in a SUMO-dependent mechanism. The truncated filaments can reanneal with another ULF to form larger VIFs. However, the phosphorylated ULF must be SUMOylated by PIAS1 to increase either the solubility or interaction with protein phosphatases, such as type-1 (PP1) and type-2A (PP2A) protein phosphatases, as shown by the large increase in vimentin phosphorylation levels with the SUMO deficient vimentin construct.^47^ These results suggest that the PIAS1-mediated SUMOylation of VIM stimulates the dephosphorylation of ULF and facilitate the reentry of the ULF into the VIF maturation process by annealing on growing VIFs. This dynamic assembly and disassembly of VIFs thus involve the SUMOylation of VIM, a modification that also regulates the cell proliferation, migration and invasion (Fig. 6).

## DISCUSSION

We report the functional effect of E3 SUMO ligase PIAS1 in HeLa cells, and determined that PIAS1 not only promotes cell proliferation, but also stimulates cell migration and invasion. PIAS1 has been extensively studied in other cancer lines, such as Human Prostate Cancer, where PIAS1 expression is increased and enhanced proliferation through inhibition of p21^21^. In addition, other studies have also reported that PIAS1 may function as a tumor suppressor to regulate gastric cancer cell metastasis by targeting the MAPK signaling pathway^51^. IL-11 mediated decrease in HTR-8/SVneo cells invasiveness was associated with a decrease in ERK1/2 activation, PIAS1/3-mediated activated STAT3 (Tyr-705) sequestration, and a decrease in PIAS1 expression, leading to a decrease in the expression of Fos and major families of metalloproteinase (MMP2, MMP3, MMP9 and MMP23B)^52^. However, these studies were limited to individual PIAS1 targets to understand the regulation mechanism. These targeted approaches sufficed to answer specific questions about PIAS1 mediated SUMOylation, but lack the depth to fully elucidate the function of PIAS1. A systematic approach to establish the global properties of PIAS1 as an E3 SUMO ligase and how these SUMOylation events alter substrate function are missing and needed. Indeed, such a method was never conceived due to the complex nature of quantitative SUMO proteomics. Global SUMO proteome analyses are challenging due to the low abundance of protein SUMOylation and the extremely large remnant that is retained on the modified lysine residues upon tryptic digestion. Moreover, proteomic workflows that are currently available to study SUMOylation require two levels of enrichment, which adversely affects the reproducibility of SUMO site quantitation.

We devised an efficient method for the identification PIAS1 substrates by modifying our previously described SUMO proteomics approach^27^. This method combines SILAC labelling for reproducible quantitative proteomic analyses, E3 SUMO ligase protein overexpression, followed by SUMO remnant immunoaffinity enrichment. This workflow allows for the selective profiling of substrates and regulated SUMOylation sites of any E3 SUMO ligase. All the PIAS1 substrates identified in this work were analyzed using forward and reverse SILAC labeling under basal condition, which further increases the confidence of the identified substrates. Notably, we observed that PIAS1 overexpression has a global effect on protein SUMOylation (Fig. 3b). This is in part due to some PIAS1 substrates being directly involved in protein SUMOylation, such as PIAS2, PIAS3, NSMCE2, TOPORS and TRIM28. In addition, many of the identified substrates were found to participate in protein ubiquitination regulation, such as TRIM24, TRIM33, RNF2 and BRCA1, which may also affect the global protein SUMOylation through the interplay between SUMOylation and ubiquitination^27, 53^.

We identified five SUMOylation sites on PIAS1 itself (Lys-46, Lys-56, Lys-137, Lys-238 and Lys-315), suggesting a possible feedback mechanism that could keep SUMOylation levels in check. Our immunofluorescence studies show that SUMOylation of PIAS1 promotes its localization to PML nuclear bodies (Figs. 5c and 5d). Interestingly, a recent paper that studied the substrates of RNF4 identified PIAS1 as a substrate of this SUMO-targeted ubiquitin ligase^54^. Moreover, RNF4 is localized to PML nuclear bodies, where it ubiquitylates SUMOylated proteins for their subsequent proteasomal degradation. Elevated levels of cellular SUMOylation may lead to an increase SUMOylation of PIAS1, prompting its localization to PML nuclear bodies and its degradation by RNF4 in a ubiquitin-dependent manner. This feedback mechanism used to regulate global SUMOylation may not be reserved solely for PIAS1. Indeed, other members of the PIAS family, as well as NSE2 and TOPORS, have been found to be SUMOylated at several lysine residues, while also being substrates of RNF4^54^.

Notably, cytoskeletal proteins occupy a significant proportion of the identified PIAS1 substrates. Constituents of actin filaments, intermediate filaments and microtubules were all found to be PIAS1 substrates. Interestingly, unlike UBC9 substrates that are typically SUMOylated on consensus motifs^6^, the acceptor lysine residues found on these cytoskeletal proteins are highly conserved but are located in non-consensus sequence motif. These observations suggest that PIAS1 may act as an adaptor protein to change cytoskeletal protein turnover or dynamics by facilitating their SUMOylation. We uncovered that SUMOylation on the tail domain of VIM increases its solubility and promotes the uptake of ULF onto VIF in a phospho-dependent mechanism. The dynamic VIF disassembly/reassembly that is promoted by VIM SUMOylation in turn favors cell motility and invasion, which could lead to an increase in cancer cell aggressiveness. Although these findings could reveal the molecular mechanism of PIAS1 mediated VIM SUMOylation and its involvement in cancer cell aggressiveness, additional evidence is required to further understand the function of PIAS1-mediated SUMOylation on the other cytoskeletal proteins and how these cytoskeletal proteins collaborate during cell migration.

## Abbreviations

SIM: SUMO-interacting motif
Ni-NTA: nickel-nitrilotriacetic acid
SCX: strong cation exchange
TCE: total cell extract
DMP: dimethyl pimelimidate
TFA: trifluoroacetic acid
ACN: acetonitrile
SUMO: small ubiquitin-related modifier
PML: Promyelocytic leukemia protein
PTM: post-translational modification
LC-MS/MS: liquid chromatography-tandem mass spectrometry
FDR: false discovery rate
PIAS1: Protein Inhibitor of Activated STAT, 1
SENPs: SUMO specific proteases
VIM: Vimentin
ULF: Unit-length filament
VIFs: Vimentin Intermediate Filaments

## ACKNOWLEDGEMENTS

This work was funded in part by the Natural Sciences and Engineering Research Council (NSERC) (P.T., RGPIN-2018-04193) and the Canadian Institute for Health Research (CIHR) (G.E., PJT 148943, PJT 148560). C.L. was supported by a scholarship from Fonds de recherche du Québec – Nature et technologies (FRQNT). The Institute for Research in Immunology and Cancer (IRIC) receives infrastructure support from Genome Canada, the Canadian Center of Excellence in Commercialization and Research, the Canadian Foundation for Innovation, and the Fonds de recherche du Québec - Santé (FRQS).

## AUTHOR CONTRIBUTIONS

C.L., F.P.M., C.P., C.M.P., T.N., L.E.A.D. performed experiments and analyzed data; C.L., F.P.M., and P.T. wrote the manuscript; G.E. and P.T. developed the concept and managed the project.

## METHODS

Methods and any associated references are available in the online version of the paper.

*Note: Supplementary information is available in the online version of the paper*

### Online Methods

#### Cell Culture and Transfection

Human cervical cancer cell line (HeLa) and HEK293 stably expressing the 6xHis-SUMO3-Q87R/Q88N mutant (HEK293-SUMO3m)^27^ were cultured in Dulbecco’s modified Eagle’s medium (HyClone) supplemented with 10% fetal bovine serum (Wisent), 1% L-glutamine (Thermo Fisher Scientific), 1% penicillin/streptomycin (Invitrogen) in 5% CO2 at 37 °C. The mammalian expression vector for Myc-PIAS1 and 6xHis-SUMO3 were constructed by inserting the full-length cDNAs into pcDNA3.0-Myc and pcDNA3.0-6xHis vectors, respectively. The mammalian expression vector for PIAS1-GFP-WT was constructed by cloning full-length PIAS1 cDNA into pcDNA3.1-c-GFP10. The PIAS1-GFP-3xKR plasmid was generated by site-directed mutagenesis using the GENEART Site-Directed Mutagenesis System according to the instructions of the manufacturer (InvitrogenTM). The mammalian expression vectors pReceiver-M11 (Flag-NSMCE2, Flag-PFDN2 and Flag-VIM) and the bacterial expression vectors pReceiver-B11 (His-NSMCE2 and His-PFDN2) were purchased from Genecopoeia, Inc. (Rockville, MD).

For transient transfection, HeLa cells or HEK293-SUMO3m cells were transfected with 1 μg plasmid per million cells using JetPrime Reagent (Polyplus-transfection) according to the manufacturer’s protocol. Cells were collected 48 h after transfection and protein overexpression was confirmed by western blot.

#### Cell Proliferation Assay

The cell proliferation assay was carried out using WST-1 Cell Proliferation Assay Kit (Roche). Transfected cells were seeded into 96-well plates at a density of 1 × 10^3^ cells/well. After 0, 2 and 4 days of incubation, WST-1 reagents were added to each well, and cells were further incubated at 37 °C for 1 h. Each measurement was performed in triplicate, and the experiments were repeated three times. The relative numbers of viable cells were estimated using the absorbance optical density (OD) at 450 nm.

#### Migration Assays

Cell migration was assessed by using a wound-healing assay. Transfected cells were seeded as a single monolayer into 12-well plates at 80%-90% confluence and then scratched using a 200-μl pipette tip. After the scratch, the cells were washed twice with PBS to remove the cell debris and grown in DMEM containing 1% FBS. Phase-contrast microscopy images of the wound area were taken at time 0 h, 24 h and 48 h, wound healing was estimated by measuring the relative width.

#### In vitro Transwell Invasion Assays

Invasive ability of transfected HeLa cells was assessed using BioCoat Matrigel Invasion Chamber (Corning) which consists of a 24-well Falcon TC Companion Plate and 8 μm pore size PET membrane with a thin layer of MATRIGEL Basement Membrane Matrix. Chambers were rehydrated by adding 0.5 ml DMEM containing 10% FBS to the interior of the inserts and bottoms of wells. The chambers were incubated in a humidified tissue culture incubator at 37°C for 2 h. After rehydration, the medium was carefully removed without disturbing the layer of Matrigel. Transfected HeLa cells were seeded at 2.5 × 10^4^ cells in 0.5 ml of serum-free DMEM to the interior of the inserts. DMEM containing 10% FBS was added to the bottoms of wells. Following a 48 h incubation at 37°C, adherent cells on the upper layer of the insert were removed by gentle scraping with a cotton tip applicator. Cells that had invaded the underside of the inserts were fixed with 100% methanol for 10 minutes at −20°C and stained with 0.5% crystal violet dye (EMD Millipore) for 10 min at room temperature. The permeated cells were imaged under a Phase-contrast microscope. Then cells were incubated with 10% acetic acid to elute the dye and the relative cell numbers were estimated using the absorbance optical density (OD) at 590 nm.

#### SILAC Labeling and Protein Extraction

HEK293-SUMO3m cells were grown in DMEM (Thermo Fisher Scientific) containing light (0Lys, 0Arg), medium (4Lys, 6Arg) or heavy (8Lys, 10Arg) isotopic forms of lysine and arginine (Silantes) for at least 6 passages to ensure full labeling. For each triple SILAC experiment, the control channel was transfected with an empty-pcDNA3.0-Myc vector while the other two channels were transfected with the Myc-PIAS1 plasmid. After 48 h transfection, an equal amount of cells from each SILAC channel were combined and washed twice with ice-cold PBS, lysed in NiNTA denaturing incubation buffer (6 M Guanidinium HCl, 100 mM NaH_2_PO_4_, 20 mM 2-Chloroacetamide, 5 mM 2-Mercaptoethanol, 10 mM Tris-HCl pH=8) and sonicated. Protein concentration was determined using micro Bradford assay (Bio-Rad).

#### Protein Purification, Digestion and Desalting

For each replicate, 16 mg of total cell extract (TCE) were incubated with 320 μL of NiNTA beads (Qiagen) at 4 °C. After 16 h incubation, NiNTA beads were washed once with 10 mL of NiNTA denaturing incubation buffer, 5 times with 10 mL of NiNTA denaturing washing buffer (8 M urea, 100 mM NaH_2_PO_4_, 20 mM imidazole, 5 mM 2-Mercaptoethanol, 20 mM Chloroacetamide, 10 mM Tris-HCl pH=6.3) and twice with 10 mL of 100 mM ammonium bicarbonate. Protein concentration was determined using micro Bradford assay (Bio-Rad). Protein digestion on beads was carried out using a ratio 1:50 sequencing grade modified trypsin (Promega): protein extract in 100 mM ammonium bicarbonate at 37 °C overnight. To quench the reaction, 0.1% trifluoroacetic acid (TFA) was added. The solution was desalted on hydrophilic-lipophilic balance (HLB) cartridges (1cc, 30 mg)(Waters) and eluted in LoBind tubes (Eppendorf) before being dried down by Speed Vac.

#### SUMO Peptide Enrichment

PureProteome protein A/G magnetic beads (Millipore) were equilibrated with anti-K(NQTGG) antibody at a ratio of 1:2 (v/w) for 1 h at 4 °C in PBS. Saturated beads were washed 3 times with 200 mM triethanolamine pH=8.3. For crosslinking, 10 μl of 5 mM DMP in 200 mM triethanolamine pH=8.3 was added per μl of slurry and incubated for 1 h at room temperature. The reaction was quenched for 30 minutes by adding 1 M Tris-HCl pH=8 to a final concentration 5 mM. Cross-linked beads were washed 3 times with ice-cold PBS and once with PBS containing 50% glycerol. The tryptic digests were resuspended in 500 ul PBS containing 50% glycerol and supplemented with cross-linked anti-K-(NQTGG) at a ratio of 1:2 (w/w). After 1 h incubation at 4 °C, anti-K-(NQTGG) antibody bound beads were washed three times with 1 ml of 1 × PBS, twice with 1 ml of 0.1 × PBS and once with ddH2O. SUMO peptides were eluted three times with 200 μl of 0.2% formic acid in water and dried down by Speed Vac.

#### SCX Fractionation

Enriched SUMO peptides were reconstituted in water containing 15% acetonitrile and 0.2% formic acid and loaded on conditioned strong cation exchange (SCX) StageTips (Thermo Fisher Scientific). Peptides were eluted with ammonium formate pulses at 50, 75, 100, 300, 600 and 1,500 mM in 15% acetonitrile, pH=3. Eluted fractions were dried down by Speed Vac and stored at −80 °C for MS analysis.

#### Mass Spectrometry Analysis

Peptides were reconstituted in water containing 0.2% formic acid and analyzed by nanoflow-LC-MS/MS using an Orbitrap Fusion Mass spectrometer (Thermo Fisher Scientific) coupled to a Proxeon Easy-nLC 1000. Samples were injected on a 300 μm ID × 5 mm trap and separated on a 150 μm × 20 cm nano-LC column (Jupiter C18, 3 μm, 300 A, Phenomenex). The separation was performed on a linear gradient from 7 to 30% acetonitrile, 0.2% formic acid over 105 minutes at 600 nl/min. Full MS scans were acquired from m/z 350 to m/z 1,500 at resolution 120,000 at m/z 200, with a target AGC of 1E6 and a maximum injection time of 200 ms. MS/MS scans were acquired in HCD mode with a normalized collision energy of 25 and resolution of 30,000 using a Top 3 s method, with a target AGC of 5E3 and a maximum injection time of 3,000 ms. The MS/MS triggering threshold was set at 1E5 and the dynamic exclusion of previously acquired precursor was enabled for 20 s within a mass range of ±0.8 Da.

#### Data Processing

MS data were analyzed using MaxQuant (version 1.5.3.8)^55, 56^. MS/MS spectra were searched against UniProt/SwissProt database (http://www.uniprot.org/) including Isoforms (released on 10 March 2015). The maximum missed cleavage sites for trypsin was set to 2. Carbamidomethylation (C) was set as a fixed modification and acetylation (Protein N term), oxidation (M), deamination (NQ) and NQTGG (K) were set as variable modifications. The option match between runs was enabled to correlate identification and quantitation results across different runs. The false discovery rate for peptide, protein, and site identification was set to 1%. SUMO sites with a localization probability of >0.75 were retained. The mass spectrometry proteomics data have been deposited to the ProteomeXchange Consortium (http://proteomecentral.proteomexchange.org) via the PRIDE partner repository with the dataset identifier <PXD011932>. Reviewer account details: Username, Password: ScCPBsuC

#### Bioinformatics Analysis

Classification of identified PIAS1 substrates was performed using PANTHER (Protein Analysis Through Evolutionary Relationships) (http://www.pantherdb.org), which classifies genes and proteins by their functions^57, 58^. The identified PIAS1 substrates were grouped into the biological process, molecular function and cellular component classes against the background of quantified SUMOylome using DAVID Bioinformatics Resources 6.7^59^. The aligned peptide sequences with ±6 amino acids sounding the modified lysine residue obtained in Andromeda were submitted to IceLogo^60^. For peptide sequences corresponding to multiple proteins, only the leading sequence was submitted. The secondary structures surrounding the PIAS1-regulated SUMO sites were investigated using NetSurfP-1.1 (Petersen et al. 2009)^61^. INTERPRO Protein Domains Analysis and Protein-Protein Interaction (PPI) network of identified PIAS1 substrates were built by searching against the STRING (Search Tool for the Retrieval of Interacting Genes/Proteins) database version 9.1^62, 63^. All predictions were based on experimental evidence with the minimal confidence score of 0.4, which is considered as the highest confidence filter in STRING. PPI networks were then visualized by Cytoscape v3.5.1^64, 65^.

#### Recombinant Protein Purification and in vitro SUMO Assay

The bacterial expression vectors pReceiver-B11 for His-NSMCE2 and His-PFDN2 recombinant protein expression were purchased from Genecopoeia, Inc (Rockville, MD). pReceiver-B11 was individually transformed into ArcticExpress Competent Cells. Transformed bacteria were cultured in LB medium until an OD600 of 0.5. Isopropyl β-D-1-thiogalactopyranoside (IPTG) (Bioshop) was added to the culture at a final concentration of 1 mM and the desired expression of the protein was inducted for 24h at 13°C. Harvested bacterial pellets were lysed and sonicated in a solution containing 500 mM NaCl, 5 mM imidazole, 5 mM β-Mercaptoethanol, 50 mM Tris-HCl pH=7.5. His-tagged proteins were purified on NiNTA beads (Qiagen). Purified proteins were eluted with 500 mM NaCl, 250 mM imidazole, 5 mM β-mercaptoethanol, 50 mM Tris-HCl pH=7.5 and concentrated on 3 kDa Ultra centrifugal filters (Amicon). Recombinant human PCNA protein was a generous gift from Dr. Alain Verreault (University of Montreal, Canada). The *in vitro* SUMO assay was carried out in a buffer containing 5 mM MgCl_2_, 5 mM ATP, 50 mM Tris-HCl pH=7.5. The enzymes responsible for the SUMOylation reaction were added at the following concentrations: 0.1 μM SAE1/2, 1 μM Ubc9, 10 μM SUMO-3 and 60 nM PIAS1. The reactions were incubated at 37 °C for 4 h and analyzed by western blot using an anti-His antibody.

#### In vivo SUMO Assay and Immunoprecipitation

HEK293-SUMO3m cells were co-transfected with Myc-PIAS1, Flag-NSMCE2, Flag-PFDN2 or Flag-VIM. At 48 h post-transfection, cells were harvested and lysed with 1 ml of Triton lysis buffer (150 mM NaCl, 0.2% Triton X-100, 1 mM EDTA, 10% Glycerol, 50 mM Tris-HCl pH 7.5) supplemented with protease inhibitor cocktail (Sigma Aldrich) at 4 °C for 15 min with gentle rocking. TCE were incubated with anti-Flag M2 Affinity Agarose Gel (Sigma Aldrich) with gentle rocking at 4 °C for overnight. Immunoprecipitates were then washed three times with cold Triton lysis buffer and were analyzed by western blot using an anti-His antibody (Sigma Aldrich).

#### Western Blot

TCE prepared in Triton lysis buffer were diluted in Laemmli buffer (10% (w/v) glycerol, 2% SDS, 10% (v/v) 2-mercaptoethanol and 62.5 mM Tris-HCl, pH=6.8), boiled for 10 min and separated on a 4–12% SDS-PAGE (Bio-rad) followed by transfer onto nitrocellulose membranes. Before blocking the membrane for 1 h with 5% non-fat milk in TBST (Tris-buffered saline with Tween 20), membranes were briefly stained with 0.1% Ponceau-S in 5% acetic acid to represent total protein content. Membranes were subsequently incubated overnight with a 1:1000 dilution of antibodies at 4 °C. Membranes were then incubated with peroxidase-conjugated anti-mouse or anti-rabbit IgG (Cell Signaling Technology) for 1 h at room temperature at a 1:5000 dilution. Membranes were washed three times with TBST for 10 minutes each and revealed using ECL (GE Healthcare) as per the manufacturer’s instructions. Chemiluminescence was captured on Blue Ray film (VWR). The following antibodies were used for western blot analyses: rabbit anti-Flag polyclonal (Sigma Aldrich), rabbit anti-PIAS1 polyclonal (Cell Signaling Technology), rabbit anti-Myc polyclonal (Cell Signaling Technology), rabbit anti-His (Sigma Aldrich), rabbit anti-Tubulin polyclonal antibody (Cell Signaling Technology).

#### Fluorescence imaging and co-localization analysis

HEK-SUMO3m cells were plated on 12-mm-diameter coverslips until they reached the desired density level and then transfected with the desired plasmids for 48 h. Cells were fixed in 4% paraformaldehyde (in PBS) for 15 min, followed by a 2-min permeabilization with 0.1% Triton X-100 (in PBS) and saturation with 2% BSA (in PBS) for 15 min. Cells were incubated with the primary antibodies for 1 h at 37 °C, rinsed and incubated with secondary antibodies conjugated to Alexa Fluor (Cell Signaling Technology) and DAPI (Sigma Aldrich) for 1 h at room temperature. Both primary and secondary antibodies are diluted in PBS / BSA 2%. To reach sub-diffraction resolution, images from fixed samples were acquired with the Airyscan detector of a Zeiss LSM880 confocal equipped with a 63X/1.43 Plan Apochromat oil immersion objective. We automatically detected and assessed the number of PIAS1 and PML positive structures by using the “particle analysis” tool from ImageJ. We then calculated the ratio manually.

#### In-gel Digestion and LC-MS/MS Identification

Cell pellets were resuspended in 5 pellet volumes of ice cold RIPA buffer. The lysate was spun at 13 000 g for 10 minutes to separate the extract into RIPA soluble and insoluble fractions. The soluble and insoluble VIM samples were separated on a 4–12% SDS-PAGE (Bio-rad), and the proteins were visualized by coomassie staining. The gel lane around vimentin corresponding position (57 kDa) was cut and then diced into ∼1 mm^3^ cubes. During the process of in-gel digestion, the gel pieces were first destained completely using destaining solution (50% H_2_O, 40% methanol, and 10% acetic acid). Then the gel pieces were dehydrated by washing several times in 50% acetonitrile (ACN) until the gel pieces shriveled and looked completely white. The proteins were reduced in 10 mM DTT at 56 °C for 30 min, alkylated in 55 mM iodoacetamide at room temperature (RT) in the dark for 30 min, and digested overnight with 300 ng of sequencing grade modified trypsin in 50 mM ammonium bicarbonate. The supernatants were transferred into Eppendorf tubes, and the gel pieces were sonicated twice in extraction buffer (67% ACN and 2.5% trifluoroacetic acid). Finally, the peptide extraction and the initial digest solution supernatant were combined and then dried using a Speed Vac. Peptides were reconstituted in water containing 0.2% formic acid and analyzed by nanoflow-LC-MS/MS using an Orbitrap Q Exactive HF Mass spectrometer (Thermo Fisher Scientific) as described previously^66^.

#### Fluorescence Recovery After Photobleaching (FRAP) assay

The FRAP assays were conducted on a LSM 880 confocal microscope equipped with a thermostatized chamber at 37 °C. The Vimentin Emerald expressing cells were detected using a GaAsp detector. Bleaching was done by combining “Time”, “Bleach” and “Region” modes on Zen software from Zeiss. Briefly, 5 pre-bleach images were taken every 5 seconds, after which five pulses of a 488 nm laser were applied to bleach an area of 25 × 2 μm. Post-bleach images were acquired every 5 seconds for 5 minutes. For fluorescence recovery analysis, the intensity in the bleached region was measured varying time points with “Frap profiler” plugin in ImageJ software. Bleach data were normalized to unbleached regions for all the time points and expressed in arbitrary units in the recovery graphs.

#### Quantification of Vimentin organization

Fixed images of cells expressing Emerald-VIM^wt^ or Emerald-VIM^mt^ were taken using a LSM 880 confocal microscope. The “title” mode in the Zen software from Zeiss was used to cover a large area of cells (2,13mm × 2,13mm). To avoid localization and conformation artefacts due to expression levels, only the cells expressing Emerald-Vimentin at an intermediate level were evaluated using the threshold module in ImageJ. The same manual threshold was used for all conditions. Cells were tabulated using the “Cell counter” plugin in ImageJ into 4 categories: ULF, VIFs, cytosolic and total cells. Categories specification were performed manually and subjectively. A representative image of each category is shown in Figure S9. More than 250 cells were counted for each experiments (N=3, n total cells = 750).

#### Statistical Analysis

Statistical analysis was carried out to assess differences between experimental groups. Statistical significance was analyzed by the Student’s t-tests. p<0.05 was considered to be statistically significant. One asterisk and two asterisks indicate p<0.05 and p <0.01, respectively.

## References

1. Flotho, A. & Melchior, F. in Annual Review of Biochemistry, Vol 82 Vol. 82 Annual Review of Biochemistry 357–385 (Annual Reviews, 2013).

2. Liang, Y. C. et al. SUMO5, a Novel Poly-SUMO Isoform, Regulates PML Nuclear Bodies. Scientific Reports 6, 15, doi:10.1038/srep26509 (2016).

3. Kunz, K., Piller, T. & Muller, S. SUMO-specific proteases and isopeptidases of the SENP family at a glance. J Cell Sci 131, doi:10.1242/jcs.211904 (2018).

4. Mukhopadhyay, D. & Dasso, M. Modification in reverse: the SUMO proteases. Trends in Biochemical Sciences 32, 286–295, doi:10.1016/j.tibs.2007.05.002 (2007).

5. Liu, B. & Shuai, K. Regulation of the sumoylation system in gene expression. Current Opinion in Cell Biology 20, 288–293, doi:10.1016/j.ceb.2008.03.014 (2008).

6. Sampson, D. A., Wang, M. & Matunis, M. J. The small ubiquitin-like modifier-1 (SUMO-1) consensus sequence mediates Ubc9 binding and is essential for SURIO-1 modification. Journal of Biological Chemistry 276, 21664–21669, doi:10.1074/jbc.M100006200 (2001).

7. Johnson, E. S. Protein modification by SUMO. Annual Review of Biochemistry 73, 355–382, doi:10.1146/annurev.biochem.73.011303.074118 (2004).

8. Shuai, K. & Liu, B. Regulation of gene-activation pathways by pias proteins in the immune system. Nature Reviews Immunology 5, 593–605, doi:10.1038/nri1667 (2005).

9. Rytinki, M. M., Kaikkonen, S., Pehkonen, P., Jaaskelainen, T. & Palvimo, J. J. PIAS proteins: pleiotropic interactors associated with SUMO. Cellular and Molecular Life Sciences 66, 3029–3041, doi:10.1007/s00018-009-0061-z (2009).

10. Tan, J. A. T., Song, J., Chen, Y. & Durrin, L. K. Phosphorylation-Dependent Interaction of SATB1 and PIAS1 Directs SUMO-Regulated Caspase Cleavage of SATB1. Molecular and Cellular Biology 30, 2823–2836, doi:10.1128/mcb.01603-09 (2010).

11. Okubo, S. et al. NMR structure of the N-terminal domain of SUMO ligase PIAS1 and its interaction with tumor suppressor p53 and A/T-rich DNA oligomers. Journal of Biological Chemistry 279, 31455–31461, doi:10.1074/jbc.M403561200 (2004).

12. Kipp, M. et al. SAF-Box, a conserved protein domain that specifically recognizes scaffold attachment region DNA. Molecular and Cellular Biology 20, 7480–7489, doi:10.1128/mcb.20.20.7480-7489.2000 (2000).

13. van den Akker, E. et al. FLI-1 functionally interacts with PIASx alpha, a member of the PIAS E3 SUMO ligase family. Journal of Biological Chemistry 280, 38035–38046, doi:10.1074/jbc.M502938200 (2005).

14. Duval, D., Duval, G., Kedinger, C., Poch, O. & Boeuf, H. The ‘PINIT’ motif, of a newly identified conserved domain of the PIAS protein family, is essential for nuclear retention of PIAS3L. FEBS Lett. 554, 111–118, doi:10.1016/s0014-5793(03)01116-5 (2003).

15. Palvimo, J. J. PIAS proteins as regulators of small ubiquitin-related modifier (SUMO) modifications and transcription. Biochemical Society Transactions 35, 1405–1408, doi:10.1042/bst0351405 (2007).

16. Kahyo, T., Nishida, T. & Yasuda, H. Involvement of PIAS1 in the sumoylation of tumor suppressor p53. Molecular Cell 8, 713–718, doi:10.1016/s1097-2765(01)00349-5 (2001).

17. Gross, M. et al. Distinct effects of PIAS proteins on androgen-mediated gene activation in prostate cancer cells. Oncogene 20, 3880–3887, doi:10.1038/sj.onc.1204489 (2001).

18. Megidish, T., Xu, J. H. & Xu, C. W. Activation of p53 by protein inhibitor of activated Stat1 (PIAS1). Journal of Biological Chemistry 277, 8255–8259, doi:10.1074/jbc.C2000001200 (2002).

19. Liu, B. et al. Inhibition of Stat1-mediated gene activation by PIAS1. Proceedings of the National Academy of Sciences of the United States of America 95, 10626–10631, doi:10.1073/pnas.95.18.10626 (1998).

20. Driscoll, J. J. et al. The sumoylation pathway is dysregulated in multiple myeloma and is associated with adverse patient outcome. Blood 115, 2827–2834, doi:10.1182/blood-2009-03-211045 (2010).

21. Hoefer, J. et al. PIAS1 Is Increased in Human Prostate Cancer and Enhances Proliferation through Inhibition of p21. American Journal of Pathology 180, 2097–2107, doi:10.1016/j.ajpath.2012.01.026 (2012).

22. Puhr, M. et al. PIAS1 is a determinant of poor survival and acts as a positive feedback regulator of AR signaling through enhanced AR stabilization in prostate cancer. Oncogene 35, 2322–2332, doi:10.1038/onc.2015.292 (2016).

23. Rabellino, A. et al. PIAS1 Promotes Lymphomagenesis through MYC Upregulation. Cell Reports 15, 2266–2278, doi:10.1016/j.celrep.2016.05.015 (2016).

24. Kadare, G. et al. PIAS1-mediated sumoylation of focal adhesion kinase activates its autophosphorylation. Journal of Biological Chemistry 278, 47434–47440, doi:10.1074/jbc.M308562200 (2003).

25. Streich, F. C. & Lima, C. D. Capturing a substrate in an activated RING E3/E2-SUMO complex. Nature 536, 304–+, doi:10.1038/nature19071 (2016).

26. Rabellino, A. et al. The SUMO E3-ligase PIAS1 Regulates the Tumor Suppressor PML and Its Oncogenic Counterpart PML-RARA. Cancer Res. 72, 2275–2284, doi:10.1158/0008-5472.can-11-3159 (2012).

27. Lamoliatte, F., McManus, F. P., Maarifi, G., Chelbi-Alix, M. K. & Thibault, P. Uncovering the SUMOylation and ubiquitylation crosstalk in human cells using sequential peptide immunopurification. Nature Communications 8, 11, doi:10.1038/ncomms14109 (2017).

28. Hendriks, I. A. et al. Uncovering global SUMOylation signaling networks in a site-specific manner. Nature Structural & Molecular Biology 21, 927–936, doi:10.1038/nsmb.2890 (2014).

29. Snider, N. T., Weerasinghe, S. V., Iniguez-Lluhi, J. A., Herrmann, H. & Omary, M. B. Keratin hypersumoylation alters filament dynamics and is a marker for human liver disease and keratin mutation. J Biol Chem 286, 2273–2284, doi:10.1074/jbc.M110.171314 (2011).

30. Goodman, S. R. Medical cell biology. (Academic Press, 2007).

31. Hochstrasser, M. SP-RING for SUMO: new functions bloom for a ubiquitin-like protein. Cell 107, 5–8 (2001).

32. Brown, J. R. et al. SUMO Ligase Protein Inhibitor of Activated STAT1 (PIAS1) Is a Constituent Promyelocytic Leukemia Nuclear Body Protein That Contributes to the Intrinsic Antiviral Immune Response to Herpes Simplex Virus 1. Journal of Virology 90, 5939–5952, doi:10.1128/jvi.00426-16 (2016).

33. Lane, E. B., Hogan, B. L. M., Kurkinen, M. & Garrels, J. I. CO-EXPRESSION OF VIMENTIN AND CYTOKERATINS IN PARIETAL ENDODERM CELLS OF EARLY MOUSE EMBRYO. Nature 303, 701–704, doi:10.1038/303701a0 (1983).

34. Ramaekers, F. C. S. et al. COEXPRESSION OF KERATIN-TYPE AND VIMENTIN-TYPE INTERMEDIATE FILAMENTS IN HUMAN METASTATIC CARCINOMA-CELLS. Proceedings of the National Academy of Sciences of the United States of America-Biological Sciences 80, 2618–2622, doi:10.1073/pnas.80.9.2618 (1983).

35. Thomas, J. T., Hubert, W. G., Ruesch, M. N. & Laimins, L. A. Human papillomavirus type 31 oncoproteins E6 and E7 are required for the maintenance of episomes during the viral life cycle in normal human keratinocytes. Proceedings of the National Academy of Sciences of the United States of America 96, 8449–8454, doi:10.1073/pnas.96.15.8449 (1999).

36. Anastasi, E. et al. Expression of Reg and cytokeratin 20 during ductal cell differentiation and proliferation in a mouse model of autoimmune diabetes. European Journal of Endocrinology 141, 644–652, doi:10.1530/eje.0.1410644 (1999).

37. Chu, Y. W., Runyan, R. B., Oshima, R. G. & Hendrix, M. J. C. EXPRESSION OF COMPLETE KERATIN FILAMENTS IN MOUSE L-CELLS AUGMENTS CELL-MIGRATION AND INVASION. Proceedings of the National Academy of Sciences of the United States of America 90, 4261–4265, doi:10.1073/pnas.90.9.4261 (1993).

38. Bordeleau, F., Bessard, J., Sheng, Y. & Marceau, N. Keratin contribution to cellular mechanical stress response at focal adhesions as assayed by laser tweezers. Biochemistry and Cell Biology-Biochimie Et Biologie Cellulaire 86, 352–359, doi:10.1139/o08-076 (2008).

39. Hu, G., Jia, F., Gao, N. & Han, Y. Impacts of CyhospitalclinE downstream vimentin on proliferation, invasion and apoptosis of hepatoma HepG2 cell. International Journal of Clinical and Experimental Medicine 11, 5564–5571 (2018).

40. Snider, N. T. & Omary, M. B. Post-translational modifications of intermediate filament proteins: mechanisms and functions. Nature Reviews Molecular Cell Biology 15, 163–177, doi:10.1038/nrm3753 (2014).

41. Zhu, Q. S. et al. Vimentin is a novel AKT1 target mediating motility and invasion. Oncogene 30, 457–470, doi:10.1038/onc.2010.421 (2011).

42. Snider, N. T., Weerasinghe, S. V. W., Iniguez-Lluhi, J. A., Herrmann, H. & Omary, M. B. Keratin Hypersumoylation Alters Filament Dynamics and Is a Marker for Human Liver Disease and Keratin Mutation. Journal of Biological Chemistry 286, 2273–2284, doi:10.1074/jbc.M110.171314 (2011).

43. Goldman, R. D., Cleland, M. M., Murthy, S. N. P., Mahammad, S. & Kuczmarski, E. R. Inroads into the structure and function of intermediate filament networks. Journal of Structural Biology 177, 14–23, doi:10.1016/j.jsb.2011.11.017 (2012).

44. Helfand, B. T. et al. Vimentin organization modulates the formation of lamellipodia. Molecular Biology of the Cell 22, 1274–1289, doi:10.1091/mbc.E10-08-0699 (2011).

45. Perez-Sala, D. et al. Vimentin filament organization and stress sensing depend on its single cysteine residue and zinc binding. Nature Communications 6, doi:10.1038/ncomms8287 (2015).

46. Satelli, A. & Li, S. L. Vimentin in cancer and its potential as a molecular target for cancer therapy. Cellular and Molecular Life Sciences 68, 3033–3046, doi:10.1007/s00018-011-0735-1 (2011).

47. Eriksson, J. E. et al. Specific in vivo phosphorylation sites determine the assembly dynamics of vimentin intermediate filaments. Journal of Cell Science 117, 919–932, doi:10.1242/jcs.00906 (2004).

48. Robert, A., Rossow, M. J., Hookway, C., Adam, S. A. & Gelfand, V. I. Vimentin filament precursors exchange subunits in an ATP-dependent manner. Proc. Natl. Acad. Sci. U. S. A. 112, E3505–E3514, doi:10.1073/pnas.1505303112 (2015).

49. Premchandar, A. et al. Structural Dynamics of the Vimentin Coiled-coil Contact Regions Involved in Filament Assembly as Revealed by Hydrogen-Deuterium Exchange. Journal of Biological Chemistry 291, 24931–24950, doi:10.1074/jbc.M116.748145 (2016).

50. Chou, Y. H., Khuon, S., Herrmann, H. & Goldman, R. D. Nestin promotes the phosphorylation-dependent disassembly of vimentin intermediate filaments during mitosis. Molecular Biology of the Cell 14, 1468–1478, doi:10.1091/mbc.E02-08-0545 (2003).

51. Chen, P. et al. Protein inhibitor of activated STAT-1 is downregulated in gastric cancer tissue and involved in cell metastasis. Oncology Reports 28, 2149–2155, doi:10.3892/or.2012.2030 (2012).

52. Suman, P. et al. AP-1 Transcription Factors, Mucin-Type Molecules and MMPs Regulate the IL-11 Mediated Invasiveness of JEG-3 and HTR-8/SVneo Trophoblastic Cells. Plos One 7, 12, doi:10.1371/journal.pone.0029745 (2012).

53. Cuijpers, S. A. G., Willemstein, E. & Vertegaal, A. C. O. Converging Small Ubiquitin-like Modifier (SUMO) and Ubiquitin Signaling: Improved Methodology Identifies Co-modified Target Proteins. Molecular & Cellular Proteomics 16, 2281–2295, doi:10.1074/mcp.TIR117.000152 (2017).

54. Kumar, R., Gonzalez-Prieto, R., Xiao, Z. Y., Verlaan-de Vries, M. & Vertegaal, A. C. O. The STUbL RNF4 regulates protein group SUMOylation by targeting the SUMO conjugation machinery. Nature Communications 8, doi:10.1038/s41467-017-01900-x (2017).

55. Cox, J. & Mann, M. MaxQuant enables high peptide identification rates, individualized p.p.b.-range mass accuracies and proteome-wide protein quantification. Nature Biotechnology 26, 1367–1372, doi:10.1038/nbt.1511 (2008).

56. Cox, J. et al. Andromeda: A Peptide Search Engine Integrated into the MaxQuant Environment. Journal of Proteome Research 10, 1794–1805, doi:10.1021/pr101065j (2011).

57. Feuermann, M., Gaudet, P., Mi, H. Y., Lewis, S. E. & Thomas, P. D. Large-scale inference of gene function through phylogenetic annotation of Gene Ontology terms: case study of the apoptosis and autophagy cellular processes. Database-the Journal of Biological Databases and Curation, 11, doi:10.1093/database/baw155 (2016).

58. Mi, H. Y. et al. PANTHER version 11: expanded annotation data from Gene Ontology and Reactome pathways, and data analysis tool enhancements. Nucleic Acids Research 45, D183–D189, doi:10.1093/nar/gkw1138 (2017).

59. Huang, D. W., Sherman, B. T. & Lempicki, R. A. Bioinformatics enrichment tools: paths toward the comprehensive functional analysis of large gene lists. Nucleic Acids Research 37, 1–13, doi:10.1093/nar/gkn923 (2009).

60. Colaert, N., Helsens, K., Martens, L., Vandekerckhove, J. & Gevaert, K. Improved visualization of protein consensus sequences by iceLogo. Nature Methods 6, 786–787, doi:10.1038/nmeth1109-786 (2009).

61. Petersen, B., Petersen, T. N., Andersen, P., Nielsen, M. & Lundegaard, C. A generic method for assignment of reliability scores applied to solvent accessibility predictions. Bmc Structural Biology 9, 10, doi:10.1186/1472-6807-9-51 (2009).

62. von Mering, C. et al. STRING: a database of predicted functional associations between proteins. Nucleic Acids Research 31, 258–261, doi:10.1093/nar/gkg034 (2003).

63. Szklarczyk, D. et al. The STRING database in 2011: functional interaction networks of proteins, globally integrated and scored. Nucleic Acids Research 39, D561–D568, doi:10.1093/nar/gkq973 (2011).

64. Shannon, P. et al. Cytoscape: A software environment for integrated models of biomolecular interaction networks. Genome Research 13, 2498–2504, doi:10.1101/gr.1239303 (2003).

65. Cline, M. S. et al. Integration of biological networks and gene expression data using Cytoscape. Nature Protocols 2, 2366–2382, doi:10.1038/nprot.2007.324 (2007).

66. McManus, F. P., Lamoliatte, F. & Thibault, P. Identification of cross talk between SUMOylation and ubiquitylation using a sequential peptide immunopurification approach. Nature protocols 12, 2342–2358, doi:10.1038/nprot.2017.105 (2017).

