## Supplementary figures for "Quantitative proteomics identifies novel PIAS1 substrates involved in cell migration and motility"

**SUMO3 (human)** N-ter…RQIRFRFDGQPINETDTPAQLEMEDEDTIDVFQQQTGG$^{92}$

**SUMO3 (mutant)** HHHHHH ..RNQTGG$^{92}$

*Figure S1: Protein sequences of the endogenous SUMO3 and SUMO3m.*

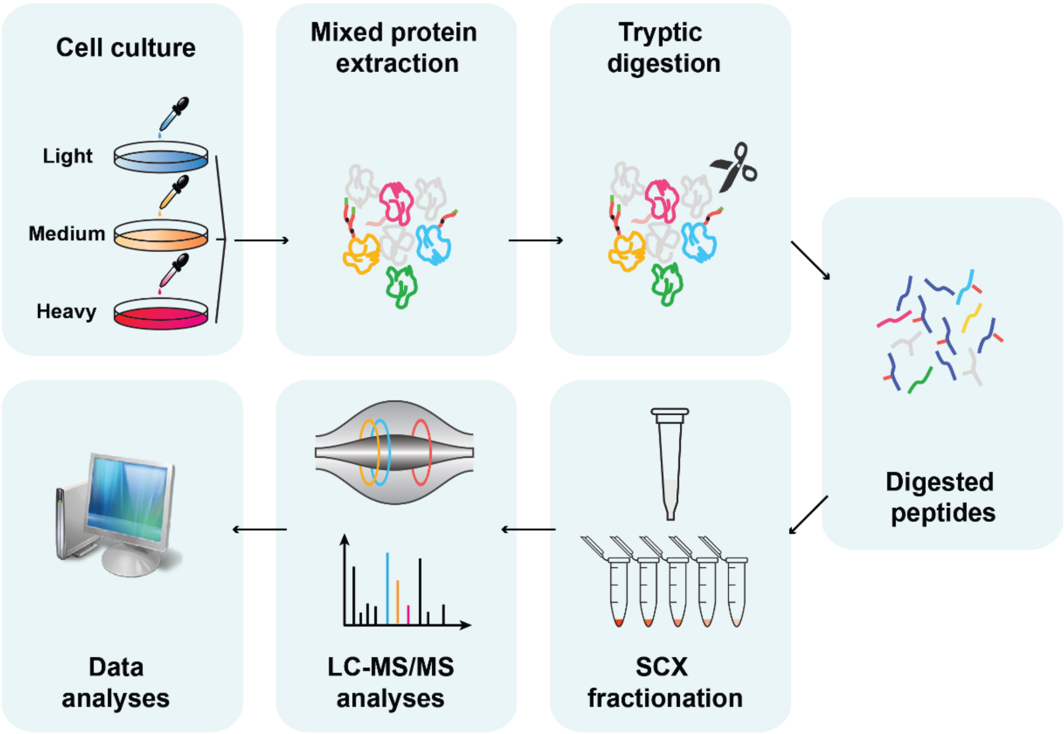


*Figure S2: Overview of proteome identification. SILAC labelled cells were lysed and combined in a 1:1:1 ratio based on protein content. Mixed cell lysates were digested using trypsin with a ratio trypsin : protein = 1:50. After desalting and drying, tryptic peptides were fractionated on SCX columns and injected on a Tribrid Fusion mass spectrometer. Peptide identification and quantification was performed using MaxQuant.*

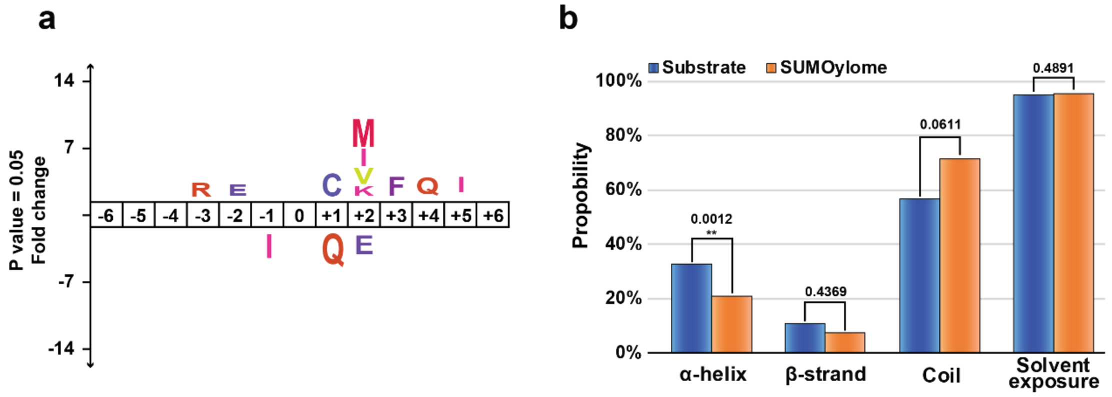


*Figure S3: Structural analysis of identified substrates.* (a) Icelogo of the amino acid sequence surrounding the PIAS regulated SUMO sites compared to the whole SUMO proteome. (b) Secondary structure predication of identified PIAS1 substrates vs identified SUMOylome.

Actin filaments

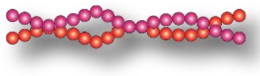

7-nm diameter

**Actin**

```
H. sapiens:      100 PEEHPVLLTEAPLNPKANREKMTQIMFETFN 128
M. musculus:     100 PEEHPVLLTEAPLNPKANREKMTQIMFETFN 128
B. taurus:       100 PEEHPVLLTEAPLNPKANREKMTQIMFETFN 128
G. gallus:       100 PEEHPVLLTEAPLNPKANREKMTQIMFETFN 128
X. tropicalis:   100 PEEHPVLLTEAPLNPKANREKMTQIMFETFN 128
D. rerio:        100 PEEHPVLLTEAPLNPKANREKMTQIMFETFN 128
                     *******************************
```

*Figure S4. Cartoon representation of the identified SUMOylation sites on Actin at Lys 115.* Protein sequence alignment of Actin across six different species showing that Lys 115 is highly conserved.

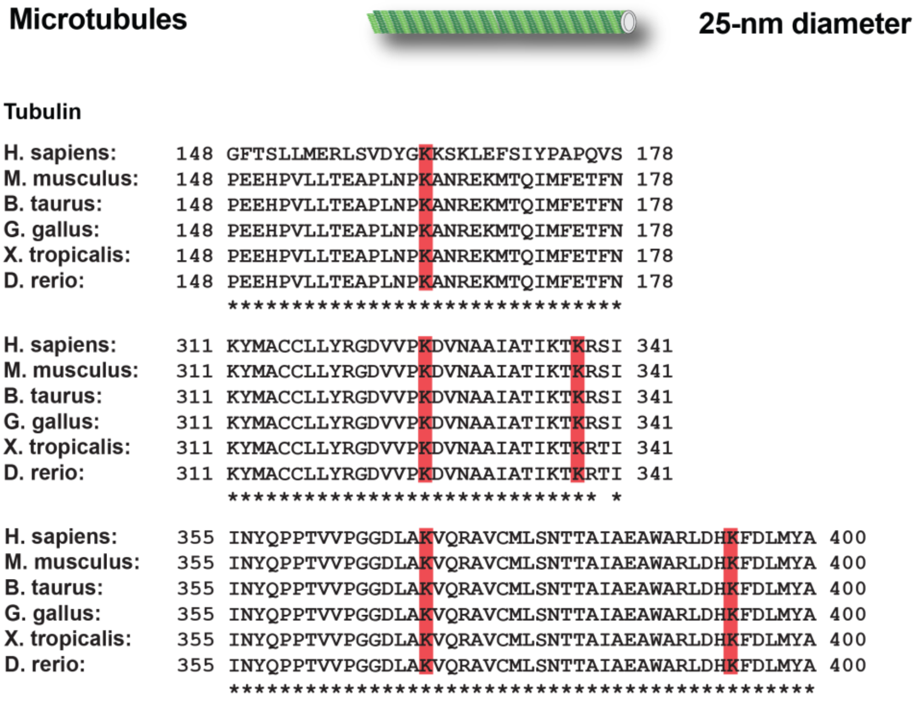


*Figure S5. Cartoon representation of the identified SUMOylation sites on Tubulin at Lys 163, Lys 326, Lys 338, Lys 370 and Lys 394.* Protein sequence alignment of Tubulin across six different species showing that all these lysines are highly conserved.

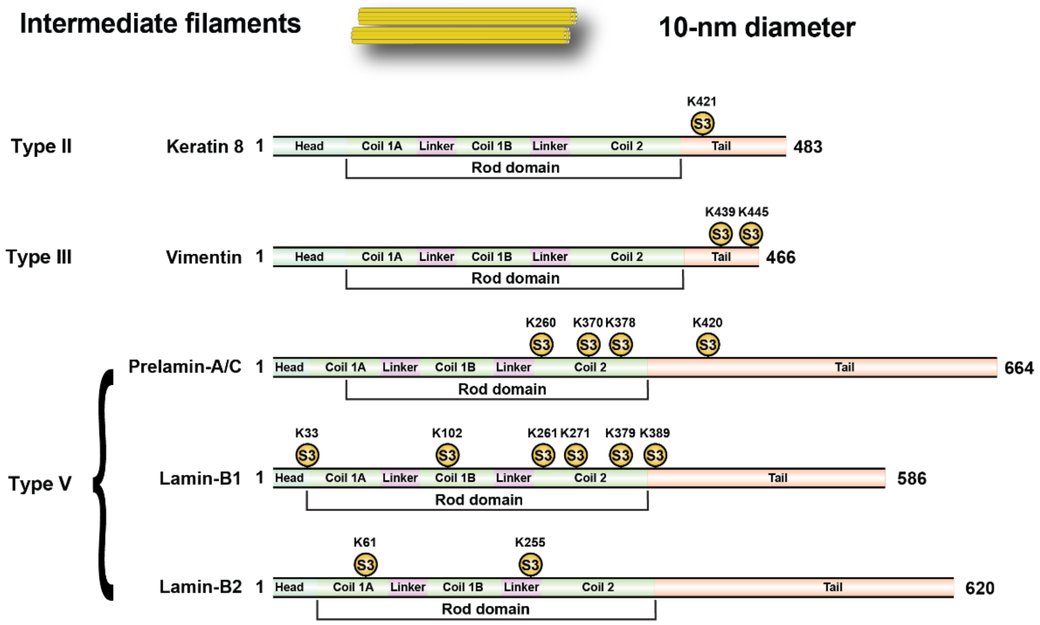


*Figure S6. Cartoon representation of the identified SUMOylation sites on different intermediate filament proteins*

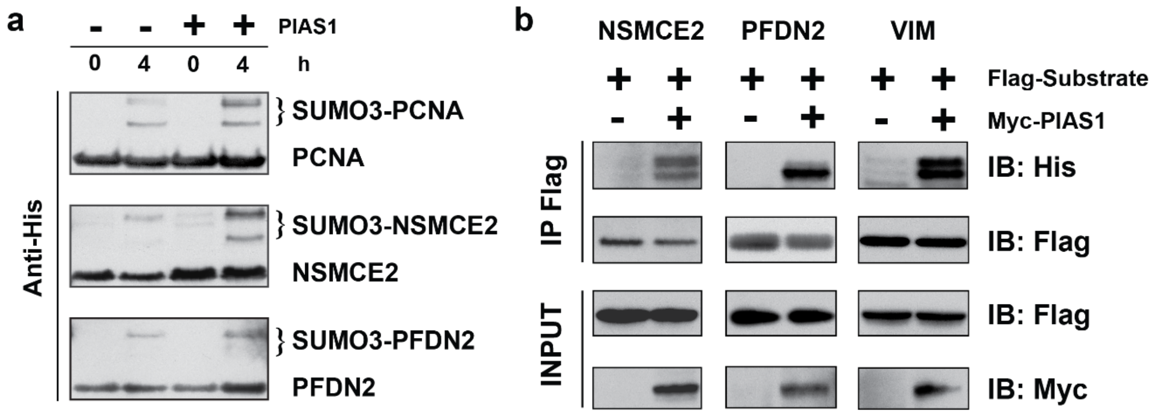


***Figure S7. Validation of SUMOylation on identified PIAS1 substrates.*** (a) *In vitro* SUMOylation assay was performed with or without PIAS1 in a buffer containing SAE1/SAE2, UBC9, SUMO3, ATP and substrates. The samples were incubated at 37 °C for 4h and examined by western blot. *In vitro* SUMO assays show that PIAS1 enhances SUMOylation of PCNA, NSMCE2 and PFDN2. (b) HEK293 SUMO3m cells were co-transfected with the indicated vectors (top), immunoprecipitated with an anti-Flag antibody, and examined by western blot. SUMOylation of NSMCE2, PFDN2 and VIM were also enhanced by PIAS1 *in vivo*.

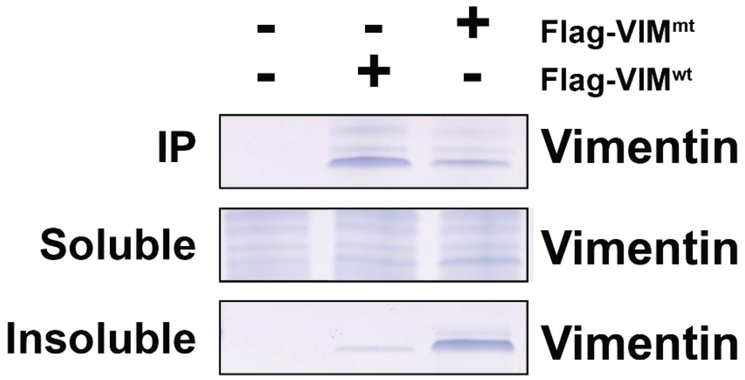


*Figure S8. SDS-PAGE gel fraction of Flag-VIM$^{wt}$, Flag-VIM$^{mt}$ and negative control from immunoprecipitation, soluble fraction and insoluble fraction used for LC-MS/MS analysis*

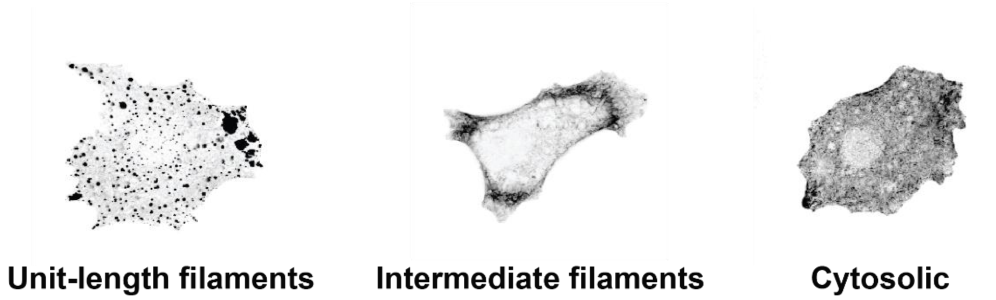


*Figure S9. Representative depiction of different forms of vimentin in MCF-7 cells.*

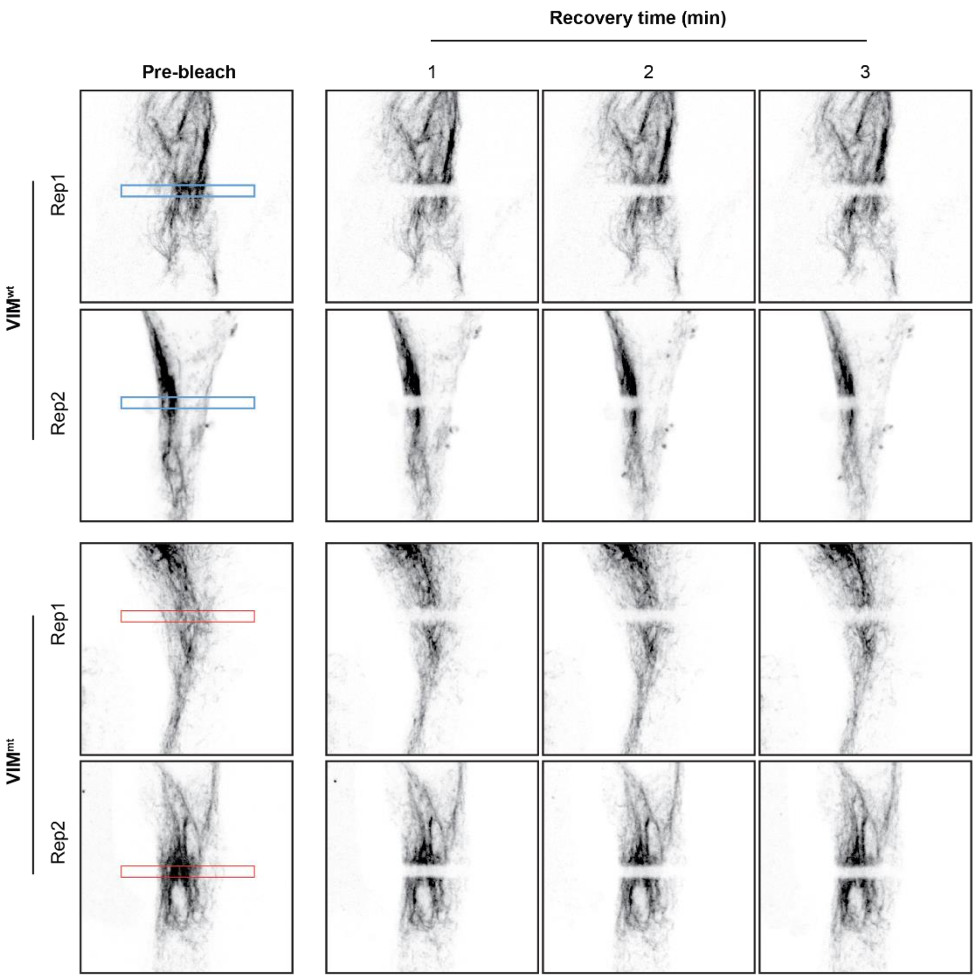


***Figure S10. FRAP assays of Emerald-VIM$^{wt}$ and VIM$^{mt}$ in MCF-7 cells.*** Selected images of fluorescence recovery after bleaching are shown.
